# Molecular Mechanism and Structural Models of Protein-Mediated Copper Transfer to the *Arabidopsis thaliana* Ethylene Receptor ETR1 at the ER Membrane

**DOI:** 10.1101/2025.01.21.634023

**Authors:** Dominik Dluhosch, Lisa Sophie Kersten, Alexander Minges, Stephan Schott-Verdugo, Holger Gohlke, Georg Groth

## Abstract

In plants, the gaseous plant hormone ethylene regulates a wide range of developmental processes and stress responses. The small unsaturated hydrocarbon is detected by a family of receptors (ETRs) located in the membrane of the endoplasmic reticulum, which rely on a monovalent copper cofactor to detect this hydrocarbon. The copper-transporting P-type ATPase RAN1 (HMA7), located in the same membrane, is known to be essential for the biogenesis of ETRs. Still, the precise molecular mechanism by which the receptors acquire their copper cofactor remains unclear. A recent study by our laboratory demonstrated a direct interaction between RAN1 and soluble copper chaperones of the ATX1 family with the model ethylene receptor ETR1, providing initial insights into the mechanism by which copper is transferred from the cytosol to the membrane-bound receptors. In this study, we further investigated these interactions with respect to the function of individual domains in complex formation. To this end, we combined biochemical experiments and computational predictions and unraveled the processes and mechanisms by which copper is transferred to ETR1 at the molecular level.

## Introduction

Ethylene is a gaseous plant hormone that plays a crucial role in a number of developmental and environmental processes in plants, including stress response, senescence, and fruit ripening. In *Arabidopsis thaliana*, the ethylene molecule is perceived by a family of membrane-bound receptor proteins (ETR1, ERS1, ETR2, EIN4, and ERS2) located at the ER membrane (Chen *et al*., 2002). All of them require an essential copper cofactor bound at their transmembrane sensor domain to sense the plant hormone with high affinity, specificity, and functionality (Rodriguez *et al*., 1999; Schaller *et al*., 1995).

The transition metal copper is an essential micronutrient for a wide range of organisms, including plants. This is largely because copper can adopt different oxidation states *in vivo*. However, this redox property contributes to its intrinsic toxicity (Festa and Thiele, 2011; Kumar *et al*., 2021; Mir *et al*., 2021; Shabbir *et al*., 2020). Redox cycling between Cu(II) and Cu(I) leads to the formation of so-called reactive oxygen species (ROS) (Kehrer, 2000), which further damage lipids, DNA, proteins, and other biomolecules (Brewer, 2010; Gill and Tuteja, 2010). Therefore, safe transportation within cells by specialized proteins is essential to ensure correct cellular functions and restrict damage induced by ROS.

In plants, several proteins have been identified that are responsible for the safe transport and modulation of intracellular copper levels. These include copper transporting transmembrane P_1B_-type heavy metal ATPases (HMAs) (Arnesano *et al*., 2002; Harrison *et al*., 1999; Harrison *et al*., 2000) and the cytosolic Cu(I)-complexing chaperones of the ATX1 family. In contrast to the cytosolic chaperones ATX1 (Andres-Colas *et al*., 2006) and CCH (Himelblau *et al*., 1998; Puig *et al*., 2007), the ER-bound transmembrane copper-transporting ATPase RAN1 has two metal binding domains (MBDs) at its N-terminus (Hirayama *et al*., 1999; Pirrung *et al*., 2008), which is distinct from the single MBD observed in the ATX1 family. Still, the precise mechanism by which copper is transported from the different plant copper chaperones to the ethylene receptors and inserted into their transmembrane domains remains largely unknown.

Our laboratory has recently uncovered a direct interaction between the transmembrane domain of *Arabidopsis* ethylene receptor ETR1 and RAN1. Furthermore, these studies provided detailed insights into the role of the soluble copper chaperones ATX1 and CCH in the intracellular transport of Cu(I) and its delivery to the receptor (Hoppen *et al*., 2019). Previous studies of HMAs and ATX1 homologs provide further insights into the copper transfer from the cytosol to the ER membrane (Andres-Colas *et al*., 2006; Hamza *et al*., 1999; Hoppen *et al*., 2019; Puig *et al*., 2007; Zhang *et al*., 2018).

To gain further insights into the molecular mechanism underlying ethylene receptor metalation, we adopted an integrated biochemical and computational approach. This allowed us to initially delineate the individual metal binding domains (MBDs) of RAN1 through computational analyses. The results were subsequently employed in biochemical, biophysical, and structural studies. To pinpoint the individual contribution of each MBD for ETR1-RAN1 complex formation, a truncated version of the receptor was employed in *in vitro* studies. This construct consisted of ETR1 domains that could only be accessed by membrane-bound RAN1, specifically the receptor’s copper-binding transmembrane and juxta-membrane GAF domains. In order to gain additional information and derive a detailed mechanism for how copper is transported from the cytosol to the ETR1 transmembrane domain, we also studied the interaction of isolated MBDs with copper chaperones ATX1 and CCH.

## Results

### Structural models and domain organization of Copper Chaperones ATX1, CCH, and RAN1

The *Arabidopsis* copper transport protein ATX1 is known to adopt a ferredoxin-like fold (*βαββαβ*-fold), a conserved structural motif observed in numerous copper-transporting proteins, including its human homolog Atox1 (Rosenzweig *et al*., 1999b). In order to emphasize the structural similarity of the selected proteins and protein domains to the ATX1 copper chaperone, we refer to this motif as the ATX1-like fold in the following. In accordance with the above, AlphaFold2 (Jumper *et al*., 2021) also predicts for ATX1 a fold consisting of the typical *βαββαβ*-fold and the “*MxCxxC*” copper-binding motif (CBM) (Rosenzweig *et al*., 1999a) (Figure S1A) (identifier: ATX1: Q94BT9) (Rosenzweig *et al*., 1999b). Similarly, CCH is known to be a homolog of ATX1 (Puig *et al*., 2007). It features the ATX1-like fold in its N-terminal metal binding domain (MBD) as well as a characteristic, extended C-terminal domain (Himelblau *et al*., 1998; Mira *et al*., 2001), as predicted by AlphaFold2 (Jumper *et al*., 2021) (identifier: CCH: O82089). A domain boundary was identified at position 72 between the MBD and the C-terminal end of CCH by the machine learning-based domain boundary prediction method TopDomain (Mulnaes *et al*., 2021b). This boundary aligns with the drop of pLDDT at position 67 in the AlphaFold model (Figure S1B) (Dluhosch *et al*., 2024; Jumper *et al*., 2021).

In contrast to soluble copper chaperones, the ER membrane-bound copper-transporting ATPase RAN1 was predicted to contain two amino-terminal metal-binding motifs, eight membrane-spanning helices, a phosphatase domain, a transduction domain, a phosphorylation domain, and an ATP-binding domain, based on the sequence similarity of RAN1 to homologous P-type copper-transporting ATPases (Hirayama *et al*., 1999) using sequence-based domain identification. This overall prediction is also reflected in the annotations in the UniProt database entry for RAN1 (identifier: RAN1: Q9S7J8). However, in contrast to the sequence-based domain identification, three domains are reported, with the last one annotated as degenerate. The AlphaFold2 model also predicts three domains at the N-terminal end, all with ATX1-like folds (henceforth referred to as MBDs) and a C_α_ atom RMSD < 1 Å with respect to each other (Figure S1C). However, the third domain (MBD3), which is the most distant from the N-terminus and the closest to the transmembrane domain, lacks the typical CBM (Figure 1A). This domain is missing in the sequence-based identification, probably due to the lack of the canonical CBM in this region. Note that a prediction with TopDomain (Mulnaes *et al*., 2021b) also identified three MBDs (Figure S1C). In bothMBD1 and MBD2, the CBM is located between the first α-helix and β-sheet of the ATX1-like fold. Additionally, unstructured regions are present at the N-terminus before MBD1 and in the linkers between the MBDs, allowing spatial flexibility of the individual domains. This is particularly relevant for MBD1, which is located most distant from the remaining RAN1 domains (Figure S1C). The boundaries predicted by TopDomain align with the observed drop in the pLDDT of the AlphaFold2 model as previously found for CCH (Figure S1C). Based on these predictions, the MBDs of the plant copper chaperons were assigned as follows: RAN1 (MBD1: AA 55-130, MBD2: AA 132-207, MBD3: AA 205-283), CCH (CCHΔ: AA 1-76, AA 77-121 were omitted) and ATX1 (ATX1: AA 1-76) (Figure S1). Assuming that copper transfer to ETR1 is achieved by direct physical interaction between the CBM of the MBDs, as proposed in a previous study (Hoppen *et al*., 2019), the individual structures of copper chaperones and MBDs were used to predict chaperone-ETR1 and chaperone-chaperone interactions, in order to elucidate copper transfer to and metalation of ETR receptors.

**Figure 1:**
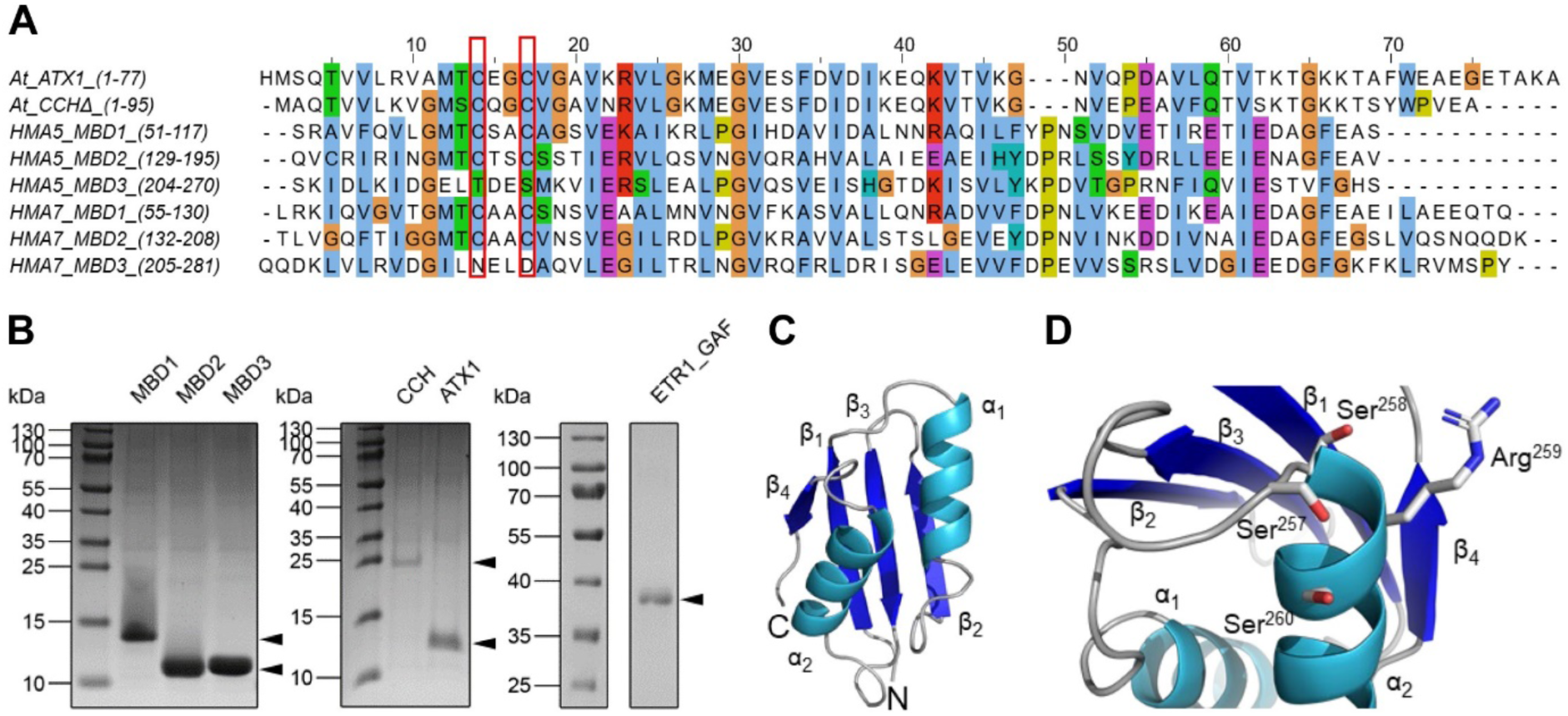
Characteristics and purity of copper chaperones, metal binding domains, and ETR1 protein constructs. **A:** Multiple sequence alignment of ATX1, the N-terminal metal binding domains (MBD) of CCH (CCHΔ), and MBDs of HMA5 and HMA7 (RAN1) from *Arabidopsis thaliana*. Amino acids are colored according to the Clustal X color scheme. Cysteines forming the copper binding motif (CBM) in ATX1, CCHΔ, and in MBD1-2 in HMA5 and HMA7 are highlighted by a red box. Note that these cysteines are missing in MBD3 of HMA5 and HMA7. The amino acids of the individual domains, which refer to their positions in the corresponding full-length proteins (given in brackets), represent the sequences of heterologously expressed and purified proteins used in this study. **B:** SDS-PAGE of all proteins used for *in vitro* experiments in this study. MBD1-3 refer to the MBDs of HMA7 as shown in (**A**). A truncation mutant of ETR1 containing the transmembrane- and GAF-domains (ETR1_GAF, aa 1-316) was used for subsequent interaction studies. **C:** Crystal structure of the RAN1 MBD3 from *Arabidopsis thaliana* at a resolution of 1.98 Å (PDB ID 8RNZ). **D:** Structure of the C-terminal region of the RAN1 MBD3 containing the *SSRS* sequence motif. Amino acid sidechains of this sequence motif are shown as sticks.

### Crystal structure of the degenerate metal binding domain MBD3

To corroborate the AlphaFold2 (Jumper *et al*., 2021; Varadi *et al*., 2022) prediction of a third ATX1-like fold domain at the N-terminus of RAN1, we expressed, purified, and analyzed this predicted domain by X-ray crystallography (Figure 1B, C). The obtained crystal of the RAN1 MBD3 from *Arabidopsis thaliana* belongs to the orthorhombic space group P2_1_22_1_ with unit cell constants *a* = 50.755 Å, *b* = 51.481 Å, *c* = 62.722 Å. The crystal structure was determined by molecular replacement (MR) using the AlphaFold2 (Jumper *et al*., 2021; Varadi *et al*., 2022) prediction of RAN1 (AlphaFold DB: Q9S7J8) cut to the sequence of MBD3 as a template. The structure was refined to a resolution of 1.98 Å with *R*_free_ = 23.5% and *R*_work_ = 19.2%, and MBD3 of RAN1 was modeled to completeness. Crystallographic data are summarized in **Fehler! Verweisquelle konnte nicht gefunden werden.**1. Compared to the primary sequence of the used construct, only the N-terminal part of the structure, consisting of the affinity-tag and a flexible linker, is absent from the deposited structure (PDB-ID: 8RNZ). Despite some inconclusive electron density in these regions, the protein backbone could not be successfully traced, probably due to the high flexibility of these regions.

The asymmetric unit comprises two molecules with an overall structural similarity (mutual RMSD = 0.355 Å; RMSDs to the AlphaFold2 model = 0.501/0.641 Å). Both molecules display the ATX1-like fold, which is typically found in copper chaperones and the regulatory domains of P_1B_-Tpye ATPases (Anastassopoulou *et al*., 2004; Banci *et al*., 2001; Gitschier *et al*., 1998; Rosenzweig *et al*., 1999a; Wernimont *et al*., 2000) confirming the AlphaFold2 structure prediction. The final structure of MBD3 is shown in Figure 1C. As previously stated, the CBM is absent. The cysteines have been substituted with asparagine and aspartate, in accordance with the results of sequence analysis by a multiple sequence alignment using Clustal Omega (Madeira *et al*., 2022) (Figure 1A). Accordingly, MBD3 is considered degenerate as classified in the UniProt database (Consortium, 2021) (UniProt-ID: Q9S7J8). Noteworthy, a structurally related *SxxS* (*SSRS*) sequence motif is present in a loop region and α-helix 2 close to the C-terminus on the opposite side of the protein (Figure 1D). Previous studies have reported that such an *SxxS* motif is capable of binding Cu(I)-ions (Pavlin *et al*., 2019; Stasser *et al*., 2005).

### Structural and functional integrity of purified proteins

Following the expression and purification of the various individual RAN1 MBDs, their biochemical and structural characteristics were evaluated to confirm that they were properly folded and functional, which was essential for subsequent interaction studies with the purified ethylene receptor ETR1. The successful purification of all proteins used in this study was demonstrated by SDS-PAGE (Figure 1B), and protein identity was confirmed by western blotting and immunodetection of the affinity tag (not shown).

First, the structural integrity of the purified MBDs was analyzed by CD-spectroscopy. Subsequently, their copper binding abilities were probed to further confirm protein function. CD-spectra of α-helical proteins are characterized by three distinctive extrema, with a maximum at 193 nm and two minima at about 208 nm and 222 nm (Holzwarth and Doty, 1965), while proteins that mainly consist of β-sheets display a maximum at 195 nm and a minimum at about 218 nm (Greenfield and Fasman, 1969). In contrast, disordered proteins are characterized by a minimum at 195 nm and overall minimal residual ellipticity above 210 nm (Venyaminov *et al*., 1993). The CD spectra of MBD1 and MBD3 display positive maxima at approximately 195 nm and two negative minima at approximately 208 nm and 222 nm, indicating the presence of α-helices in these MBDs. MBD2 displays the same characteristics, with a negative residual ellipticity minimum at about 222 nm, which is shifted towards 218 nm, and a positive maximum, which is shifted towards 193 nm. These differences are indicative of a higher proportion of β-sheets in MBD2 compared to MBD1 and MBD3 (Figure 2A). Overall, these data indicate the presence of secondary structure and suggest the structural integrity of the purified recombinant MBDs. Quantitative analysis by the BeStSel webserver (Micsonai *et al*., 2022; Micsonai *et al*., 2018; Micsonai *et al*., 2015) confirms the presence of secondary structure and overall structural similarity of the three RAN1 MBDs, while also indicating some minor differences in the secondary structure, particularly in the α-helix and β-sheet regions (Table S2). Note that estimates of the secondary structure composition from CD spectra may fail to provide acceptable results on α/β-mixed or β structure-rich proteins; the BeStSel method used here has been developed to mitigate such challenges (Micsonai *et al*., 2015).

**Figure 2:**
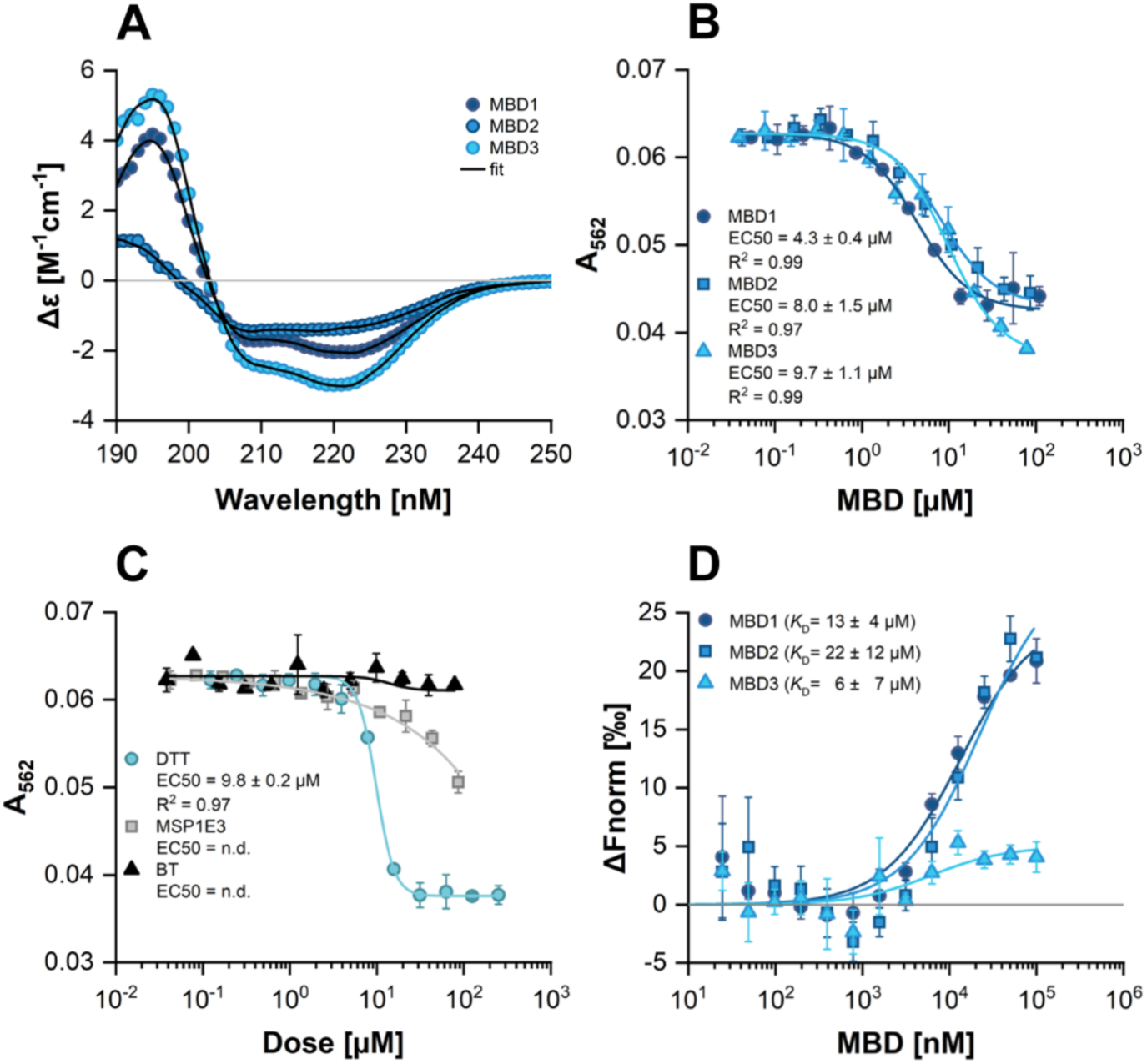
Biophysical and biochemical characterization of the RAN1 metal binding domains (MBDs) 1-3 from *Arabidopsis thaliana.* **A:** Far-UV CD-spectra of MBDs 1 to 3. **B:** Copper transfer from the chromophoric Cu(I)-BCA_2_ complex to MBDs 1 to 3 as a function of protein concentration. Competition for copper is monitored at 562 nm. **C:** Control samples used in the copper binding assay. DTT (positive control), MSP1E3 (His-tag control), and bleed-through (BT) control testing for unremoved DTT in MBD samples. Data from **B** and **C** were fit to a four-parameter logistic function and EC50-values are reported in the figure. **D:** Interaction of RAN1 MBDs 1 to 3 with ETR1_GAF^1-316^-studied by microscale thermophoresis (MST). ETR1_GAF^1-316^ is the labeled protein. Data were fit to a one-site binding model and the obtained dissociation constants (*K*_D_) are reported in the figure. In all subfigures, symbols represent experimental data and solid lines represent fitted data. All measurements were performed at least in triplicate.

Following CD-spectroscopy, we studied copper binding to the purified MBDs to further probe their structural integrity and function. To this end, a spectrophotometric assay was employed, in which the high-affinity Cu(I)-specific ligand bicinchoninic acid (BCA) competes for copper binding with other Cu(I) chelators (Dluhosch *et al*., 2024; Schott-Verdugo *et al*., 2019). In this assay, the formation of the chromophoric Cu(I)-BCA_2_ complex is reduced accordingly in the presence of copper binding proteins and indicated by a decrease in absorption. The results of this assay (Figure 2B) confirm copper binding to MBD1 and MBD2. MBD1 showed a higher efficiency to compete with BCA for copper than MBD2, as evidenced by the related EC_50_ values (EC_50 *MBD1*_ = 4.3 ± 0.4 µM and EC_50 *MBD2*_ = 8.0 ± 1.5 µM). Notably, MBD3 also showed copper-binding, with an EC_50 *MBD3*_ = 9.7 ± 1.1 µM (Figure 2B). The apparent dissociation constants *K*_D *MBD-Cu(I)*_ were computed according to (eq. 9): MBD1: 4.78 ⨉ 10^-14^ M; MBD2: 8.89 ⨉ 10^-14^ M; MBD3: 1.07 ⨉ 10^-13^ M. To eliminate the possibility that the copper binding observed with MBD3 is related to incomplete removal and carry-over of the copper chelator DTT from protein purification, buffer controls were treated the same way as MBD3 samples. However, no decrease in absorption and thereby no evidence of copper binding was observed in these samples (Figure 2C), indicating successful buffer exchange after purification prior to copper binding studies. Moreover, DTT has distinct copper binding characteristics and parameters compared to those observed in the MBD binding studies. This is evidenced by the Hill coefficient of the related binding curves (4.2 for DTT vs 1.4 for MBDs). This further substantiates that the copper binding observed with MBD3 is not caused by DTT. Further tests using a His-tag control protein (MSP1E3) indicate some extend of copper binding by the tag at high protein concentration (Figure 2C). However, in comparison to the MBDs, the observed decrease in absorption is less pronounced, and the overall curve shape does not align with the binding characteristics and parameters observed with the MBDs. Therefore, the copper binding observed with purified MBD3 cannot or at least not entirely be attributed to the His-tag, but reflects binding at MBD3.

### Interaction of soluble MBDs with ETR1

Previously published results demonstrated a direct interaction between the ETR1 transmembrane domain (ETR1_TMD) and the copper transporter RAN1 (Hoppen *et al*., 2019). Further analysis of this interaction using RAN1 subdomains revealed that the N-terminal domain containing the MBDs (NterRAN1) is directly involved in the interaction with ETR1_TMD (Hoppen *et al*., 2019). However, the localization of three MBDs in NterRAN1, of which MBD3 does not contain the typical *CxxC* binding motif, raises the question of whether all MBDs bind to ETR1 with similar affinities or if there is a preferred interaction. This, in turn, could provide novel information regarding the sequence and precise mechanism by which copper is transferred from RAN1 to ETR1. To gain detailed insight into the role of each MBD for copper transfer to the receptor in the extra-, juxta-, and intramembrane domain, we applied a receptor construct containing the TMD and the adjacent extramembrane GAF domain (ETR1_GAF) and studied the interactions between individual MBDs and this ETR1 construct. First, we performed *in vitro* interaction studies using microscale thermophoresis (MST) on ETR1_GAF and MBDs in accordance with the experiments outlined in (Hoppen *et al*., 2019). The results show that MBD1 and MBD2 bind to ETR1_GAF, with overall very similar and almost identical binding curves and apparent dissociation constants (*K*_D_) of 13.3 ± 4.4 µM for MBD1 and 22.3 ± 12.1 µM for MBD2 (Figure 2D). The situation is less clear for the interaction of the ETR1_GAF module and the degenerate metal binding domain MBD3, which shows a less pronounced signal amplitude than observed with MBD1-2 (Figure 2D). Fitting of these data to a single-site binding model, as for MBD1-2, results in an apparent *K*_D_ of 6.0 ± 7.3 µM for MDB3. Overall, the binding studies on the individual MBDs suggest that all MBDs bind to the ETR1-GAF receptor construct with similar affinities.

### Structural models of chaperone-ETR1 interactions

We used ColabFold 1.5.2 to generate structural models of the chaperone/MBD-ETR1 interactions (Mirdita *et al*., 2021). However, models obtained by this approach did not agree with experimental evidence (Hoppen *et al*., 2019) as either no interaction or only interactions with the TMD of ETR1 were predicted. Such interactions are not plausible as neither the soluble chaperones ATX1 and CCH nor the MBDs of RAN1 contain amino acids that would allow penetration of the ER membrane to reach the copper binding active site located at the middle of the bilayer. Therefore, possible interaction sites between the soluble chaperones ATX1, CCHΔ, and MBD1-3 with ETR1 were first identified using GLINTER (Graph Learning of INTER-protein contacts) (Xie and Xu, 2022). GLINTER is a deep learning method for interfacial contact predictions of dimers and is based on coevolutionary data obtained by multiple sequence alignments and structural data and can predict possible interaction sites in protein-protein complexes (Xie and Xu, 2022). The predictions obtained using this approach suggest that ATX1 and CCHΔ interact via their copper binding motif (CBM) with the GAF-domain of ETR1 (Figure 3). The construct ETR1^306-738^ used in ref. (Hoppen *et al*., 2019) lacks the GAF-domain and does not show binding of ATX1 and CCHΔ. An additional contact site is proposed for residues at the beginning of the second β-sheet of the chaperones with the ETR1 linker region connecting the transmembrane domain to the GAF-domain and with the GAF-domain itself. MBD1 and MBD2 of RAN1, like ATX1 and CCH, show interactions around their CBMs with the GAF-domain of ETR1 and interactions with the ETR1 linker region but lack contacts between their second β-sheets and the corresponding ETR1 counterparts. MBD3 is predicted to bind with the second β-sheet to the ETR1 linker region and the GAF domain, similar to ATX1 and CCH. Additional interactions of the degenerative CBM with the ETR1 linker region and GAF domain are indicated. No interaction at the alternative CBM is suggested. Interestingly, out of all tested chaperone-like domains, the predicted interactions for MBD3 with ETR1 are most confident (Figure 3), suggesting a particularly conserved interaction of MBD3 with ETR1. Note that no interactions between ETR1 and the MBDs 1 to 2 at the β-sheet were predicted, contrary to the prediction for ATX1 and CCH. In turn, MBD3 shows the most conserved interactions via the β-sheet (Figure 3).

**Figure 3:**
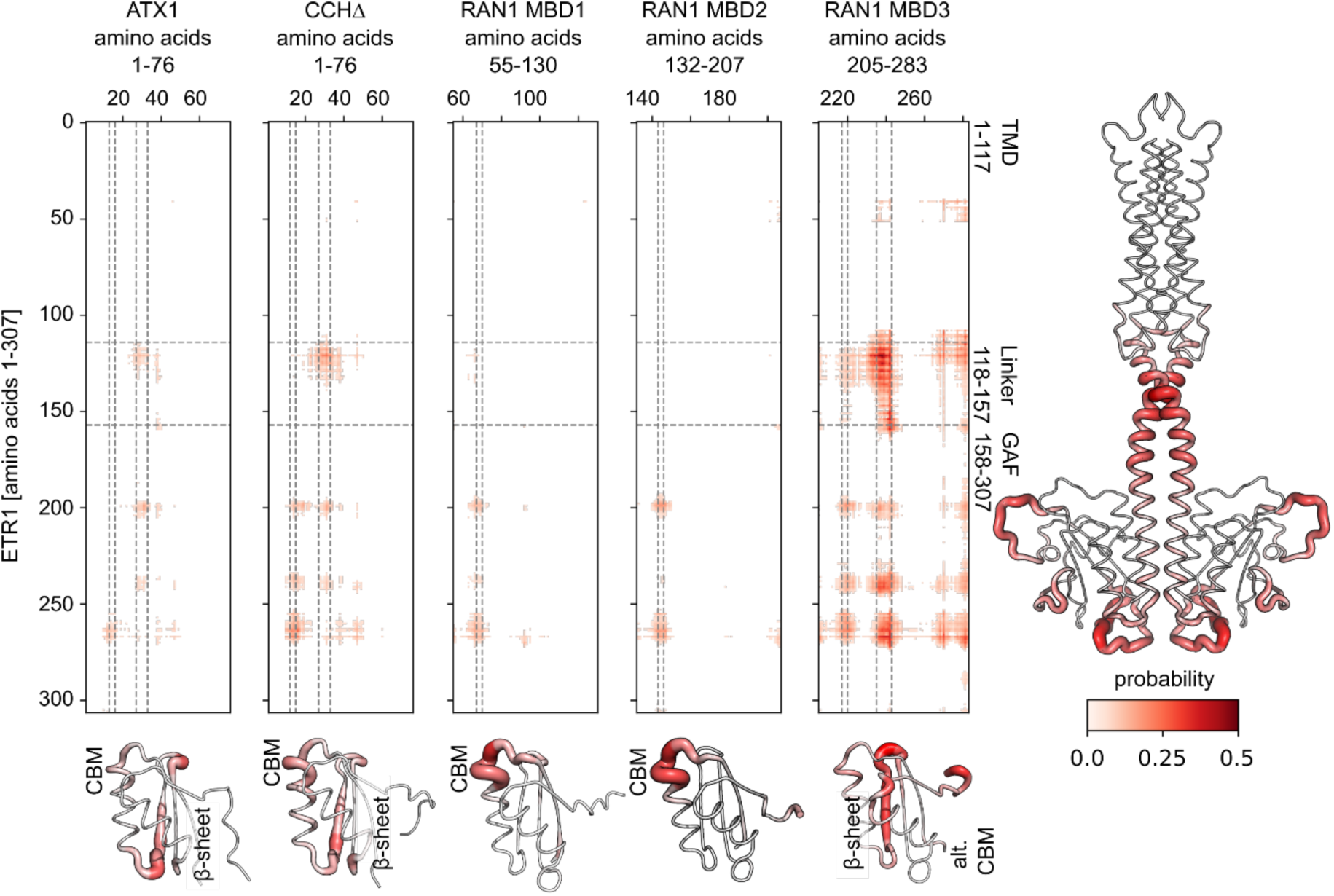
Contact maps of soluble chaperones ATX1, CCHΔ, and MBD1-3 of RAN1 with ETR1 predicted with GLINTER. The darker red color and thicker lines in the putty representation of ETR1_GAF indicate a higher probability of interactions with the chaperones or MBDs. ATX1 and CCHβ, as well as MBD1 and MBD2 of RAN1, show interactions at the CBM (vertical lines at residues 13 and 16, 67 and 70, 144 and 147, respectively) with the linker region and GAF-domain of ETR1. Additionally, interactions are proposed for the residues at the beginning of the second β-sheet (vertical lines at residues 27 and 33, 235 and 243, respectively) with the ETR1 linker region and GAF domain. The latter interactions are not present for MBD1 and MBD2. For MBD3, no interaction at the degenerate CBM is indicated. Overall, MBD3 displays a similar pattern to ATX1 and CCH, but with higher prediction confidence, showing interactions of the second β-sheet with the ETR1 linker region and GAF domain.

The predicted protein-protein contacts were further used as ambiguous restraints for protein-protein docking using HADDOCK2.4 (Dominguez *et al*., 2003). For both the full-length *ab initio* (Schott-Verdugo *et al*., 2019) (Figure 4A-E) and the AlphaFold2 ETR1-model (Figure 4F-J), the density representations of all chaperone poses around ETR1 were investigated and agreed with the defined restraints. The best-scored structures show that ATX1, CCH, and MBD3 interact with the ETR1 GAF domain (Figure 4A, B, E, F, G, J), whereas MBD1 interacts with both the linker region of ETR1 and parts of the GAF-domain (Figure 4C, H). MBD2 interacts with the linker region of ETR1 only (Figure 4D, I). The proposed interaction of MBD2 with the receiver domain of ETR1 (Figure 4D, I) is not meaningful in the biological context, given that the RAN1 copper transporter is located in the same membrane as the ETR1 receptor protein. In contrast to isolated MBD2 used in our prediction, MBD2 in RAN is too distant to access the C-terminal receiver domain of ETR1.

**Figure 4:**
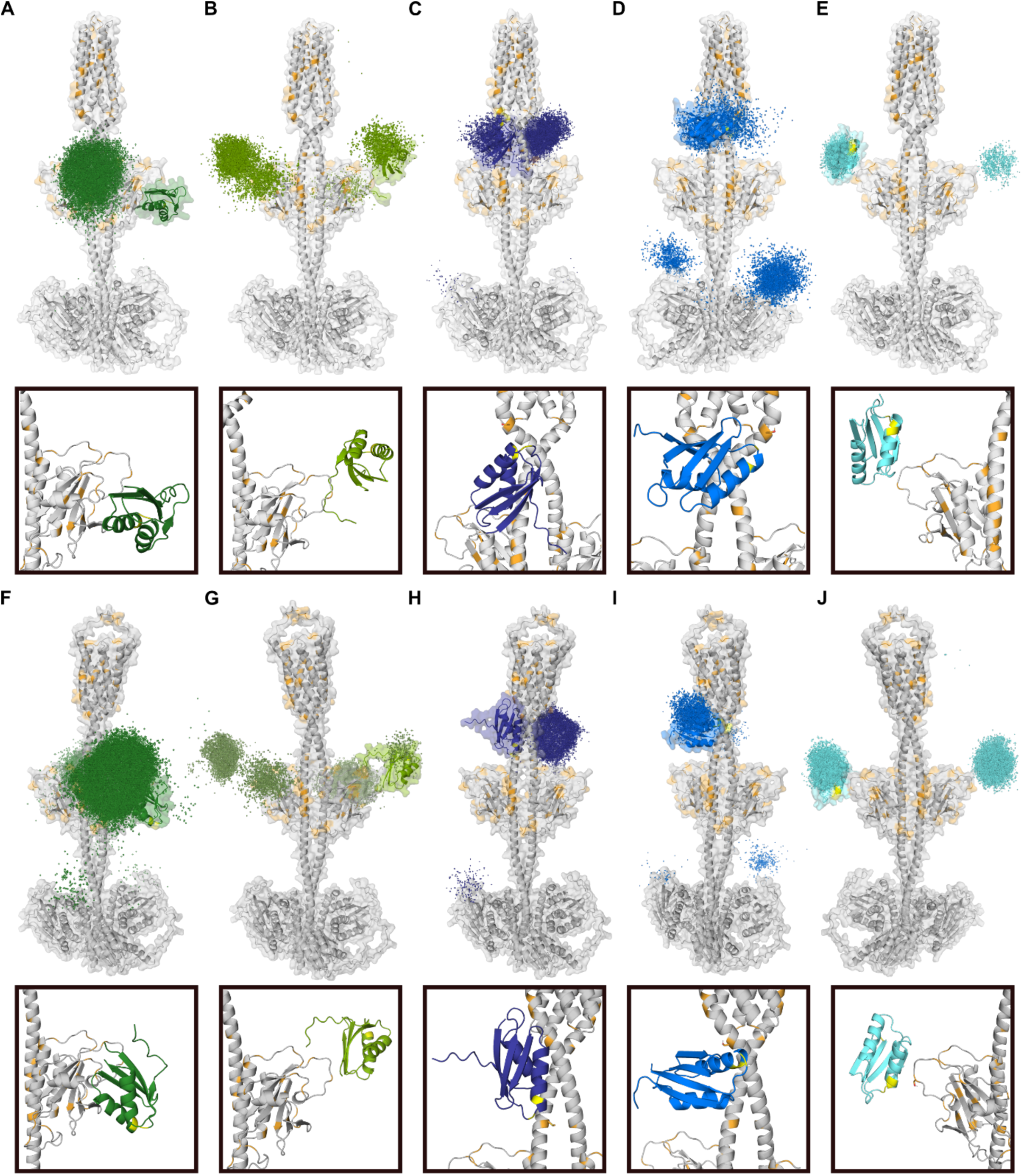
Density representation of all chaperone docking poses with ETR1 and best-scoring complexes. **A-E:** full-length *ab initio* ETR1. **F-J:** full-length ColabFold ETR1. Density representation of all ATX1 (A, F: dark green), CCH (B, G: yellow-green), MBD1 (C, H: purple), MBD2 (D, I: blue), and MBD3 (E, J: cyan) poses with ETR1 (gray) and the accordingly selected best-scored complex with the corresponding zoom. Amino acids forming the CBM in the chaperone are colored yellow. Residues of ETR1 able to complex Cu(I)-ions, such as cysteine, histidine, methionine, and serine are colored orange.

As anticipated from the contact maps used for the restraints in protein-protein docking (Figure 4), the predicted complexes of ATX1 and CCH demonstrate that the CBM is not oriented towards the TMD of ETR1. In contrast, this was observed for MBD1 and MBD2. The CBM is frequently located in close proximity to ETR1 (< 5 Å) and to residues able to interact with monovalent copper ions, such as cysteine, histidine, methionine, and serine (Rubino and Franz, 2012; Stasser *et al*., 2005). This would facilitate direct copper transfer from MBD1 and MBD2 to ETR1. In the case of MBD3, no clear orientation of the degenerate CBM could be determined.

To obtain full-length models of the ETR1-RAN1 complex, a multimeric docking attempt was performed. The MBD1 of RAN1 is expected to be the most mobile MBD given that it is the first N-terminal MBD, has the longest linker of about 8 aa to MBD2, and, unlike MBD2 and MBD3, does not interact with the transmembrane domains of RAN1 in the AlphaFold structure. To mimic this enhanced mobility of MBD1 in the rigid docking approach, MBD1 itself and a truncated version of RAN1 lacking MBD1 were docked to ETR1 collectively. As in the preceding docking approach, the predictions of GLINTER were used as ambiguous restraints to guide the docking process. Moreover, unambiguous restraints were employed to represent the loop between MBD1 and MBD2, while allowing for their movement with respect to each other. The best-scored ETR1-RAN1 complexes obtained for both full-length ETR1 models demonstrate that even in the absence of membrane restraints, a spatial arrangement was identified in which the TMDs of ETR1 and RAN1 are located in the same membrane plane (Figure 5A, D). As observed in the aforementioned docking approach (Figure 4D, E, I, J), MBD2 and MBD3 bind to the same domains (Figure 5B, E). This suggests that the interactions predicted by GLINTER and those generated by HADDOCK in the context of a full-length ETR1-RAN1 complex are consistent. In contrast to the aforementioned docking approach, MBD1 is now located at the GAF domain of ETR1 (Figure 5B, E). Given that MBD2 is already situated at the linker region of ETR1, and that the loop between MBD1 and MBD2 is too short to accommodate the binding of MBD1 on the opposite side, an alternative interaction site in the vicinity of the other monomer’s GAF-domain is proposed for MBD1 (Figure 5C), This is in line with the interaction observed for the soluble chaperones ATX1 and CCH (Figure 4A, B). The density representations of the MBDs across all ETR1-RAN1 complexes show a consistent pattern in both models, with MBD1 binding to the GAF domain, MBD2 to the linker region, and MBD3 to the side of the GAF domain (Figure 5C, F). An alternative hypothesis is that MBD1 and MBD2 bind sequentially, which would allow binding to similar regions.

**Figure 5:**
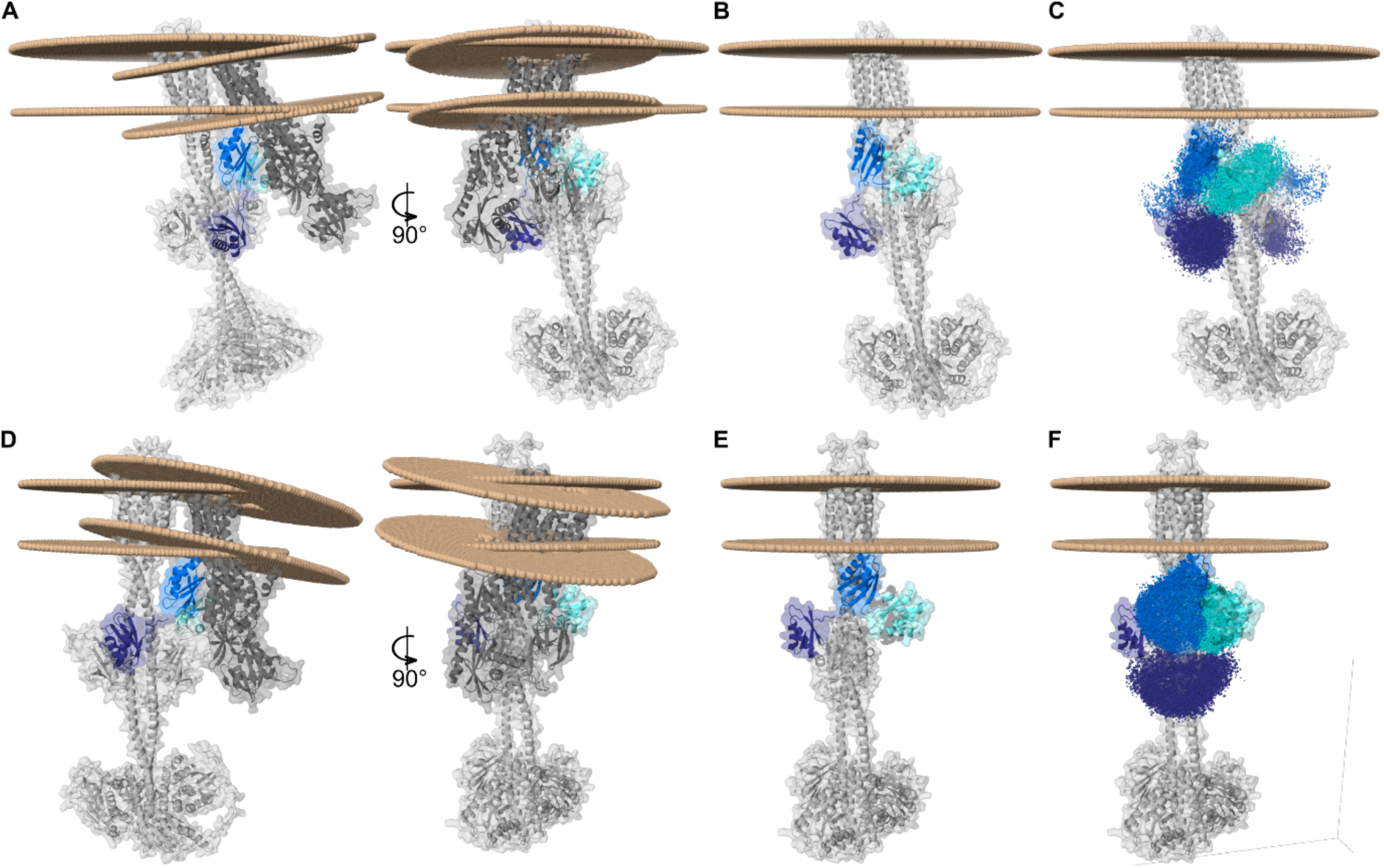
Model of the ETR1-RAN1 complex. The models indicate binding positions of the individual MBDs as obtained in previous results (Figure 3, 4) and the orientation of RAN1 and ETR1 in relation to the membrane. The three MBDs are colored purple (MBD1), blue (MBD2), and cyan (MBD3). An alternative interaction site of MBD1 in the vicinity of the GAF domain is predicted. MBD2 and MBD3 bind to the same domains as in the initial docking approach. **A:** Membrane orientation of RAN1 and full-length *ab initio* ETR1 after docking. **B:** Truncated RAN1 lacking MBD1 and MBD1 were collectively docked to full-length *ab initio* ETR1. The remaining domains of RAN1 are hidden for the sake of clarity. **C:** Density representation of MBD1, MBD2, and MBD3 poses at full-length *ab initio* ETR1 **D:** Membrane orientation of RAN1 and full-length ColabFold ETR1 after docking. **E:** Truncated RAN1 and MBD1 docked to full-length ColabFold ETR1. The remaining domains of RAN1 are hidden for the sake of clarity. **F:** Density representation of MBD1, MBD2, and MBD3 poses at full-length ColabFold ETR1.

The recently published AlphaFold3 has demonstrated substantial improvements in accuracy over many previous protein complex prediction tools (Abramson *et al*., 2024). The new AlphaFold version predicts an ETR1-RAN1 complex with transmembrane domains in a similar orientation. However, the transmembrane domains of ETR1 and RAN1 are arranged such that they would not be located in the same membrane plane (Figure S2C). As with the HADDOCK results, MBD3 of RAN1 is placed at the GAF domain of ETR1, with MBD2 positioned directly at the linker region (Figure S2D). Similar to the HADDOCK results, MBD1 is positioned between the GAF domain and the ETR1 linker region on the opposite side of ETR1, with the CBM oriented towards the GAF domain (Figure S2C). Overall, the local interactions predicted by AlphaFold3 are consistent with the aforementioned protein-protein docking results.

### Interaction of RAN1 MBDs with soluble copper chaperones ATX1 and CCH

Next, we studied the interactions of individual RAN1 MBDs with soluble copper chaperones ATX1 and CCH. This was to determine whether there is a preference for an MBD to bind to one of these chaperones and whether this binding is more likely than binding to ETR1. All MBD/chaperone combinations showed binding, yet overall, ATX1 was clearly preferred to CCH, as evidenced by the 10-20 fold higher affinities of MBD1 and MBD2 and a similar affinity of MBD3 for ATX1 in comparison to CCH. The affinity differences observed here with the single MBDs are higher than those found in ref. (Hoppen *et al*., 2019) with the N-terminal RAN1 tail containing all three MBDs, which might indicate additional steric influences when all three MBDs are present. The apparent *K*_D_-values are 154 ± 53.7 nM (ATX1-MBD1), 238.8 ± 86.8 nM (ATX1-MBD2), 105.3 ± 78.9 nM (ATX1-MBD3) versus 3164.7 ± 683.6 nM (CCH-MBD1), 2778.3 ± 989.9 nM (CCH-MBD2) and 88.5 ± 78.2 nM (CCH-MBD3) (Figure 6A, B).

**Figure 6:**
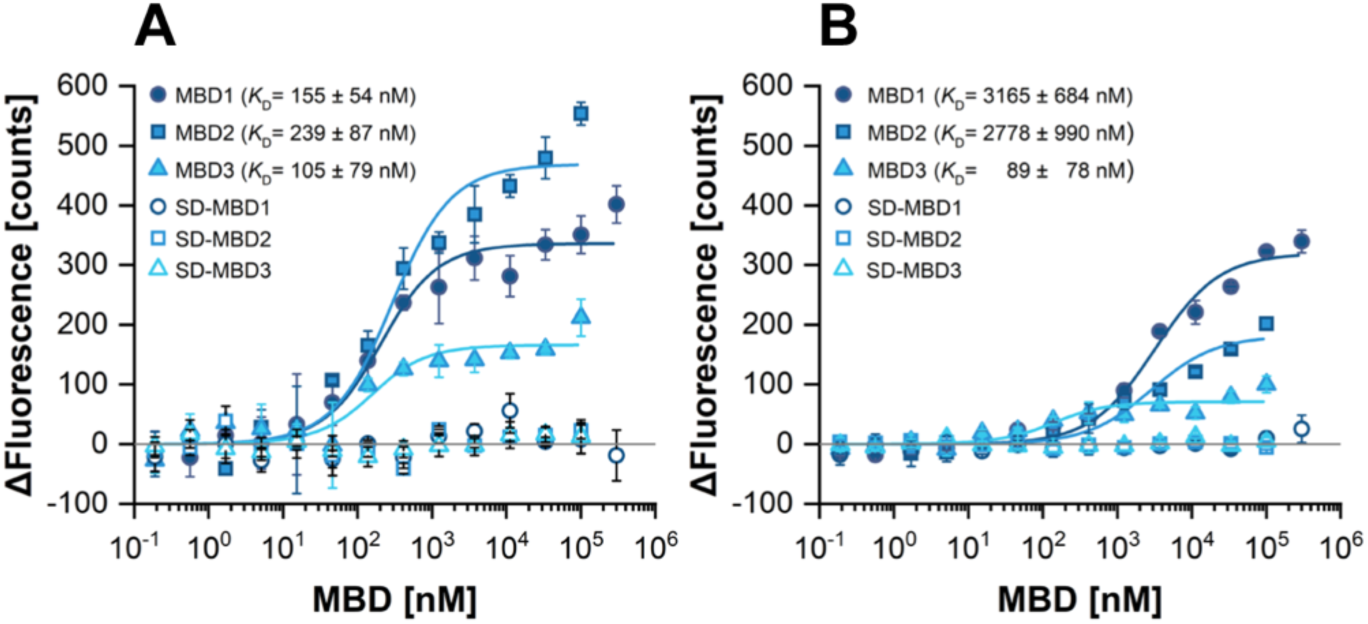
Interaction of RAN1 MBDs 1 to 3 with the soluble copper chaperones ATX1 and CCH studied by microscale thermophoresis (MST). **A:** ATX1 and **B:** CCH are the labeled proteins. Data were fit to a one-site binding model and the obtained dissociation constants (*K*_D_) are reported in the figure. In all subfigures, symbols represent experimental data and solid lines represent fitted data. All measurements were performed at least in triplicate.

The binding studies indicate that ATX1 is the preferred interaction partner of the RAN1 MBDs, putatively due to the shielding effect of the additional C-terminal domain in CCH for MBD1 and MBD2 (Himelblau *et al*., 1998). Note that binding of ATX1, CCH, and CCHΛ− to the full-length RAN1 is ∼4-5 times stronger than to the N-terminal tail of RAN1 (Hoppen *et al*., 2019) or the single MBDs (Figure 6). This observation may be attributed to the fact that the degrees of freedom in the unbound state are more restricted when the MBDs are attached to the remainder of RAN1 but binding of copper chaperones to the ATPasés translocon is also a plausible explanation since such an interaction provides an additional binding site and has been described before (Gonzalez-Guerrero and Arguello, 2008; Padilla-Benavides *et al*., 2013).

In light of the HADDOCK results, the copper relay hypothesis, and the experimentally determined interaction of MBDs with soluble chaperones, structural models of heterodimers of ATX1, CCHΔ, and MBDs were predicted using ColabFold 1.5.2 (Mirdita *et al*., 2021) (Figure 6 A to D). All individual monomers display the characteristic ATX1-like fold. The proposed heterodimers of ATX1, CCH, MBD1, and MBD2 are oriented in a face-to-face arrangement (Keller *et al*., 2012). This is in contrast to the heterodimers that have been predicted for MBD3. ATX1 and CCH form either face-to-face (Keller *et al*., 2012) or face-to-back dimers (Keller *et al*., 2012) with MBD3. MBD1 and MBD2 form back-to-back dimers exclusively with MBD3, which prevents the CBMs from approaching the other domain. As the complexes generated by ColabFold do not take into account the fact that MBD1 and MBD2 are connected by a linker of approximately eight residues, MODELLER 10.5 (Eswar *et al*., 2006) was employed to generate a complete model (Figure 7E). This model underlines that a direct copper transfer from the soluble chaperones ATX1 and CCH to MBD1, and subsequently from MBD1 to MBD2, is feasible from the structural perspective (Figure 7), which is further supported by known structural arrangements of homologous complexes (Banci *et al*., 2006; Rodriguez-Granillo and Wittung-Stafshede, 2008).

**Figure 7:**
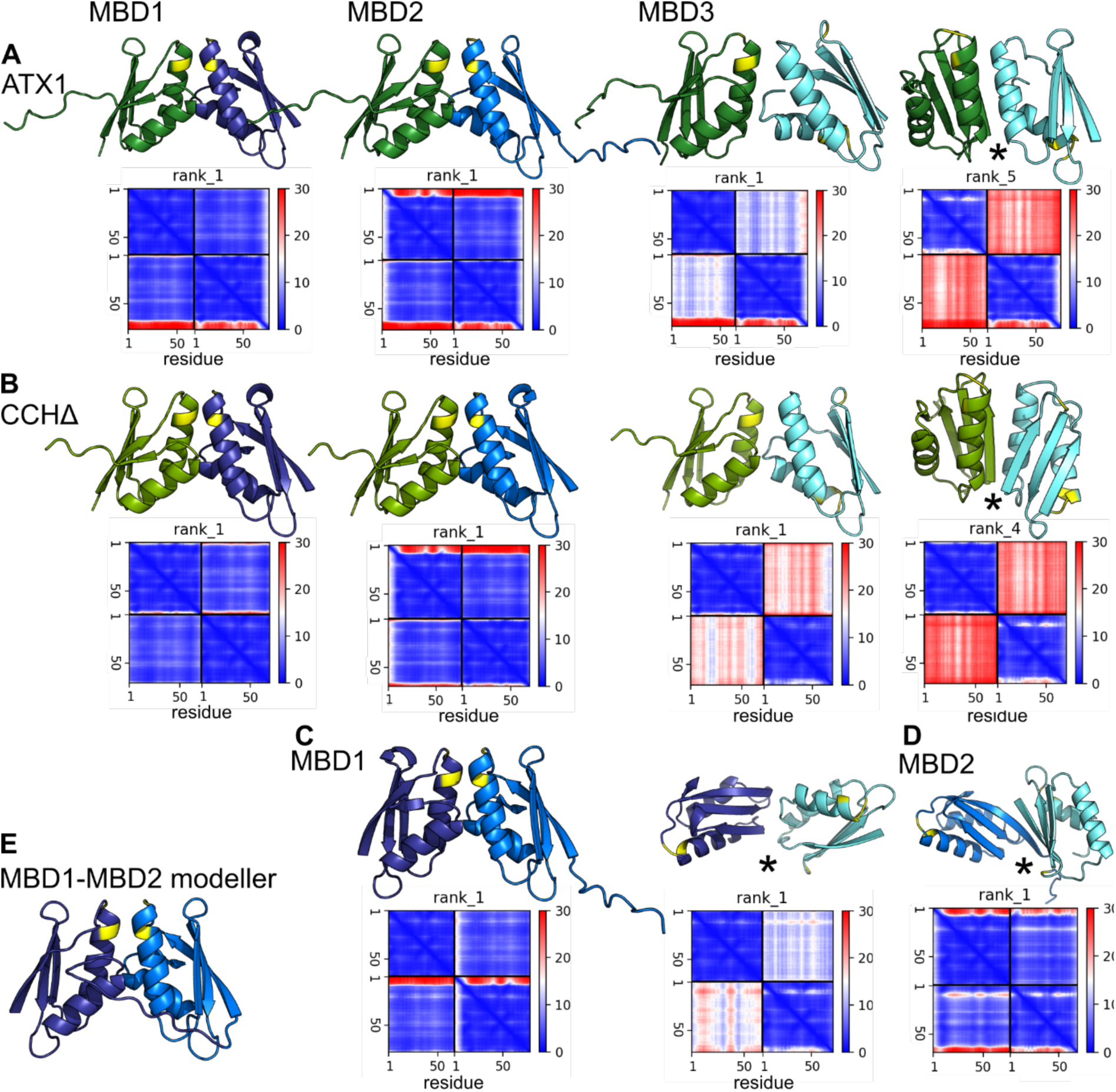
Putative models of the individual copper chaperones ATX1, CCHΔ, MBD1, MBD2, and MBD3. (ATX1: dark green, CCHΔ: light green, MBD1: purple, MBD2: blue, MBD3: cyan) and corresponding PAE (predicted alignment error matrix). All individual monomers have the typical ATX1-like fold. The heterodimers of ATX1, CCH, MBD1, and MBD2 orient in a face-to-face conformation. The soluble chaperones ATX1 and CCH from face-to-face and back-to-face dimers with MBD3. MBDs 1 and 2 form back-to-back dimers with MBD3. **A:** Proposed ATX1-MBD heterodimers. **B:** Proposed CCH-MBD heterodimers. **C:** Proposed MBD1-MBD heterodimers. **D:** Proposed MBD2-MBD3 heterodimer. **E:** MBD1-MBD2 heterodimer built based on the ColabFold prediction using MODELLER 10.5. Copper binding motifs are colored yellow.

## Discussion

Previous research demonstrated potential direct interactions between the soluble copper chaperones ATX1 and CCH and the ER-bound copper-transporting P-type ATPase RAN1 with the ethylene receptor ETR1 (Hoppen *et al*., 2019). This provided the first insights at the molecular level into receptor metalation, which is required for the sensing of and the signaling and plant response to the ethylene plant hormone. In the present study, we further dissected these interactions and propose possible structural models of copper chaperones ATX1, CCH, full-length RAN1, and individual RAN1 MBDs with the ethylene receptor ETR1 aiming to gain further insight into the sequence and molecular mechanisms underlying receptor metalation.

AI-driven methods for predicting protein structures, like AlphaFold, RoseTTAFold, and D-I-TASSER, offer unparalleled insights into the structures of individual proteins and their multimeric complexes. In this work, the AlphaFold2 structure prediction of RAN1 was corroborated by a sequence-based analysis of RAN1, which identified two MBDs in the N-terminal region. Notably, AlphaFold2 predicted a third soluble domain adopting the ATX1-like fold (therefore termed MBD3), which was not predicted by the sequence-based domain identification, likely due to the absence of a CBM in this domain. In contrast, the domain boundary prediction method TopDomain (Mulnaes *et al*., 2021b) also identified three MBD (Figure S1C). The ATX1-like fold and the existence of a third domain (MBD3) were confirmed experimentally by X-ray crystallography. The obtained crystal structure of MBD3 (Figure 1C) is consistent with the structure predicted by AlphaFold2 (C_α_ atom RMSD < 1 Å). Consequently, all three MBD domains were utilized for the structural prediction of ETR1-RAN1 and MBD-chaperone complexes.

First, GLINTER was used to predict possible interactions or contact sites between soluble chaperones ATX1 and CCH or the MBDs of RAN1 with ETR1. The data indicate that the interaction interfaces among the three MBDs are separated and that MBD3 is more likely to interact with ETR1 than ATX1 and CCH, particularly with the receptoŕs GAF domain (Figure 3). Subsequent docking experiments using the full-length forms of ETR1 and RAN1 revealed that MBD3 is positioned at the ETR1 GAF domain (Figures 5). This suggests that this interaction is also feasible when the structural characteristics and restraints of ETR1 and RAN1 are taken into account. Further support for our model that RAN1 engages in complex formation with ETR1 through its MBDs is provided by previous studies on P-type ATPases, which have demonstrated that their complex formation, e.g., in the context of dimerization, is facilitated by their cytosolic domains (Zeinert *et al*., 2024)

Further computational studies in this work on potential dimeric interactions between the MBDs or with the soluble copper chaperones revealed that heterodimers involving MBD3 assemble face-to-face but also face-to-back or back-to-back (Figure 7). Naturally, direct copper transfer between MBD3 and MBD1 or MBD2 would not be feasible in such assemblies, although the degenerate MBD3 is still capable of copper binding, as demonstrated by our copper binding studies (Figure 2B). Hence, MBD3 in RAN1 may serve a different function such as a regulatory role as described for other MBDs (Bitter *et al*., 2022; Guo *et al*., 2024) However, this does not preclude the aforementioned hypothesis that MBD3 may serve as an anchor in RAN1-ETR1 complex formation, particularly when considering our findings regarding contact site predictions through GLINTER, which makes use of coevolutionary information (Xie and Xu, 2022) (Figure 3).

Due to the low signal-to-noise ratio, the *K*_D_ of the MBD3-ETR1 interaction calculated from the *in vitro* binding studies shows a higher relative variance than for the interaction of MBD1 and MBD2 with the receptor (Figure 2D). Nevertheless, the data still suggest a somewhat higher affinity for the interaction of MBD3 with ETR1 than for the other MBDs, which is in line with the previously proposed anchor function of MBD3 in ETR1-RAN1 complex formation. The comparable affinities of MBD1 and MBD2 (*K*_D_ = 13 ± 4 µM and 22 ± 12 µM) might suggest that both MBDs bind to overlapping sites on ETR1. This hypothesis is further supported by the high structural and sequence similarity of the MBDs. Furthermore, this is in line with our docking experiments with the individual MBDs. Our results indicate that MBD1 and MBD2 are located at the ETR1 linker region, in close proximity to the ETR1 TMD where the copper cofactor is bound (Figure 4 and 5), and they do this via interactions around their CBM. Once loaded with copper, the MBDs could transfer copper ions to the receptor, suggesting a mechanism by which the receptor receives its copper cofactor. Previous copper loading of the MBDs, on the other hand, can be achieved by a copper transfer from soluble chaperones ATX1 or CCH through direct interaction. To this end, our *in vitro* interaction studies demonstrated that the binding of ATX1 to MBDs 1 to 3 is favored over the binding of MBDs to CCH (Figure 6A, B). This is consistent with the previous hypothesis that ATX1 and CCH fulfill different tasks, given that they are inversely regulated under conditions of excess copper and copper deficiency (del Pozo *et al*., 2010). ATX1 is regarded as the primary factor in intracellular copper transport, while CCH is supposed to be predominantly involved in intercellular copper transport facilitating the remobilization of copper from decaying plant organs (Mira *et al*., 2001). The proposed copper transfer mechanism is in accordance with our *in vitro* experimental data on MBD-chaperone formation (Figure 6A, B) and is further corroborated by the published literature on MBD-chaperone interaction (Arnesano *et al*., 2002; Pufahl *et al*., 1997; Walker *et al*., 2002; Wernimont *et al*., 2000).

The copper binding assay shown in Figure 2 indicates that MBD1 and MBD2 similarly compete for copper, as evidenced by the obtained apparent *K*_D_ values (4.78 ⨉ 10^-14^ M; 8.89 ⨉ 10^-14^ M). As the thermodynamic driving force due to the difference in the two *K*_D_ values is negligible, Cu(I) will likely be transferred *in vivo* to the most readily accessible MBD, which is MBD1 in the structural context of RAN1 and more specifically the RAN1-ETR1 complex (Figure 5C, F). Our docking studies identified two different interaction sites of MBD1 with ETR1. These include the linker region, which also encompasses the MBD2 binding site (Figure 4), and the GAF domain of ETR1, which is also recognized by ATX1 and CCH (Figure 4, 5). Furthermore, the study of heterodimers comprising MBD1 and either ATX1 or CCH indicates that these heterodimers adopt a face-to-face conformation, thereby enabling a direct copper transfer. Hence, MBD1 should be able to accept copper from the soluble chaperones ATX1 and CCH, although binding of CCH may be less likely due to the lower affinity for MBD1. The apparent dissociation constant of Cu(I) from ATX1 is reported to be ∼10^-18^ M (Xiao *et al*., 2011) and therefore lower than the apparent *K*_D_’s of Cu(I) from MBD1 and MBD2 determined in our study. Thus, there is no thermodynamic driving force for transferring Cu(I) from ATX1 to the MBDs. It was suggested in a similar context that copper trafficking proteins overcome the extraordinary copper chelation capacity of the cytoplasm by catalyzing the *rate* of copper transfer between physiological partners (Huffman and O’Halloran, 2000). Hence, the metallochaperones could work like “enzymes”, carefully tailoring energetic barriers along specific reaction pathways but not others.

The results of our docking experiments indicate that MBD2 is located in close proximity to the linker region and the TMD of ETR1 (Figure 2 and 3). Furthermore, computational predictions on dimeric interactions among the MBDs and with soluble copper chaperones indicate that heterodimers of MBD1 and MBD2 adopt a face-to-face conformation. Hence, a direct copper transfer seems feasible, given the proximity of the CBMs. This is supported by the *K*_D_ values of MBD1 or MBD2 for Cu(I), which are ∼40 to 80-fold higher than the dissociation constant of copper from the ETR1_TMD (Schott-Verdugo *et al*., 2019), which would provide a thermodynamic driving force for the Cu(I) transfer to the TMD of ETR1. In ref. (Zimmermann *et al*., 2009), an apparent dissociation constant of *K*_D_ of 7.94 ⨉ 10^-19^ M was reported for MBD1 (termed HMA7n there), where BCS was used as a probe in the competition rather than BCA. Further differences may arise because the apparent dissociation constants of Cu(I)-transporting systems are pH-dependent (Xiao *et al*., 2011). In light of the aforementioned results, and given the proximity of MBD2 to the TMD of ETR1, it seems plausible to suggest that MBD1 initially accepts copper from ATX1 and subsequently transfers this ion to MBD2, which then transfers it to ETR1. The RAN1 MBDs all have comparable affinities for ATX1, with an about 100-fold higher binding affinity than that observed for ETR1, thereby favoring the interaction with ATX1. Although the affinity of the individual MBDs for ETR1 is lower than for ATX1, the persistent presence of the MBDs at the membrane and their proximity to ETR1 permit a subsequent transfer of the copper ion, particularly when the ETR1-RAN1 complex is formed.

The capacity of the degenerate MBD3 to bind monovalent copper was unexpected, given the established characteristics of a canonical degenerate MBD. However, further analysis revealed that this may be related to a solvent-accessible methionine in the MBD3 C-terminus that may form a Cu(I)-binding complex with a second methionine from an additional MBD3 monomer. Yet, such an assembly would involve a dimer complex that differs substantially from known structures of related copper chaperones such as Atox1 (Levy *et al*., 2017; Perkal *et al*., 2020). Nevertheless, previous studies have demonstrated that closely spaced methionines are capable of binding monovalent copper. This includes a conserved *MxxxM* motif within the transmembrane domain of yeast copper transporter Ctr3, which was identified as essential for copper transport across membranes (Puig *et al*., 2002). Furthermore, it has been suggested that the copper transporter Ctr1 contains at least two methionines that coordinate a Cu(I)-ion within its pore (Ren *et al*., 2019). Alternatively, the unexpected copper binding of MBD3 may be related to an *SSRS* sequence motif in its C-terminus that is also solvent-exposed (Figure 1D). *In vitro* and *in silico* experiments have demonstrated that such sequence motifs are capable of binding Cu(I) ions (Pavlin *et al*., 2019; Stasser *et al*., 2005). Nevertheless, the physiological relevance of this *SxxS* sequence motif remains to be elucidated as previously outlined by Pavlin *et al*. (2019) (Pavlin *et al*., 2019). This may be achieved, for instance, through the analysis of *Cys-to-Ser* mutations, which are commonly used in analyzing the function of individual MBDs in P_1B_-Type ATPases (Yatsunyk and Rosenzweig, 2007) and in investigating Cu(I):protein ratios of copper chaperones and enzymes (Heaton *et al*., 2000; Stasser *et al*., 2005). Finally, the copper affinity of MBD3 may be an effect of the isolated domain and not be present or relevant in the *in vivo* context if there is no transfer possibility from the soluble chaperones to MBD3 for steric reasons.

In conclusion, our data suggest that the individual MBDs in RAN1 may fulfill distinct roles in receptor complex formation in addition to regulating RAN1 (Figure S3). MBD3 may function as an anchor, stabilizing the RAN1-ETR1 complex while MBD1 and MBD2 are involved in a copper relay system. In this process, MBD1 is suggested to accept the copper ion from the soluble copper chaperone ATX1 (Figure S3, step 1.) and subsequently transfer it to MBD2 (Figure S3, step 2.). Given its proximity to the ETR1 TMD, MBD2 might transfer the copper ion to the receptor (Figure S3, step 3.). Similar findings have been observed in the yeast homolog Ccc2, where the first MBD has been characterized as an acceptor domain (Banci *et al*., 2007). The hypothesis of a direct copper transfer from RAN1 to ETR1 via protein-protein interaction is supported by the previously identified direct interaction of ETR1 and RAN1, which was studied by fluorescence complementation in tobacco plants (Hoppen *et al*., 2019), as well as the described increase in ethylene binding by ETR1-containing, isolated yeast membranes when CuSO_4_ is added (Binder *et al*., 2007). These experimental findings collectively indicate that a membrane-bound protein mediates the transfer of copper to the receptor.

## Methods

### Structural models of copper chaperones ATX1, CCH and RAN1 MBDs

To date, no experimental structures of the copper chaperones ATX1, CCH, and RAN1 from *Arabidopsis thaliana* have been reported. Computational structural models are available within the AlphaFold database (identifiers: ATX1: Q94BT9, CCH: O82089, RAN1: Q9S7J8) (Jumper *et al*., 2021). The structures of homologs from yeast, humans, and *Archaeoglobus fulgidus*, among others, have been resolved (sequence identity 43-32%, 40-25%, and 48-42%, ATX1, CCH, and RAN1, respectively). To differentiate the metal binding domains (MBDs) from unstructured parts or movable linkers, *TopDomain* from *TopSuite* (Mulnaes *et al*., 2021b; Mulnaes *et al*., 2021a) was used to perform a domain boundary prediction for CCH and RAN1. The predicted boundaries were applied to the corresponding AlphaFold2 structures of ATX1, CCH, and RAN1, to extract the metal-binding domains only. The obtained MBDs of ATX1, CCH (CCHΔ), and RAN1, were then used to predict putative chaperone-chaperone and chaperone-ETR1 interactions.

### Prediction of structural models of chaperone-ETR1 interactions

Structural models of the chaperone-ETR1 interactions were generated using ColabFold 1.5.2 (Mirdita *et al*., 2021). However, the results were inconsistent with the available experimental evidence (Hoppen *et al*., 2019), as they either indicated no interaction or suggested interactions with the TMD of ETR1. Accordingly, we investigated potential interactions between the soluble copper chaperones ATX1, CCHΔ, MBD1-3, and ETR1. To this end, we employed *GLINTER* (Graph Learning of INTER-protein contacts), a deep learning method for predicting interfacial contacts of dimers (Xie and Xu, 2022). The GLINTER predictions are based on the PDB structures of the soluble copper chaperones (ATX1: aa 1-76, truncated CCH (CCHΔ): aa 1-76; aa 77-121 were omitted) as well as the MBDs of RAN1 (MBD1: aa 55-130, MBD2: aa 132-207, MBD3: aa 205-283), and the *AlphaFold2* monomer structure of ETR1 (aa 1-307). We restricted ETR1 to residues 1-307 as previous *in vitro* binding studies have demonstrated that ATX1, CCH, and CCHΔ bind to ETR1 residues 1-157, but not to ETR1 residues 306-738. With regard to the N-terminal end of RAN1, stronger interactions were measured with residues 1-157 in comparison to residues 306-738 (Hoppen *et al*., 2019). Consequently, the GAF domain of ETR1 (aa 158-306) was included, as this domain may represent a structural element that should be taken into account during docking. The twenty highest-ranked GLINTER-predicted interaction contacts were employed as restraints for protein-protein docking using HADDOCK2.4 (Dominguez *et al*., 2003). At present, two computational models of the transmembrane domain of ETR1 are available (Jumper *et al*., 2021; Schott-Verdugo *et al*., 2019; Varadi *et al*., 2022). It cannot be excluded that the discrepancies in the spatial orientations of the transmembrane helices and the connecting loops that are partially exposed to the cytoplasm may have an impact on the docking results. Consequently, a dimeric full-length ETR1 model featuring the transmembrane domain model proposed by AlphaFold2 was generated using ColabFold. Conversely, a dimeric full-length ETR1 model featuring the ab initio transmembrane model was merged with the cytosolic part of the ColabFold model using MODELLER 10.1. Prior to docking, the full-length ETR1 dimers were protonated using PROPKA3 (Olsson *et al*., 2011) in Maestro (2023-2, 2023) with pH 7.4. The obtained protonation states were used to mimic physiological conditions and were incorporated into the docking using HADDOCK2.4 (Dominguez *et al*., 2003). The protein-protein contacts predicted by GLINTER were used as ambiguous restraints with distances of 2 to 20 Å. Initial interaction models of ATX1, CCHΔ, and MBD1-3 of RAN1 with the two ETR1 models were obtained. Given that MBD1-3 are domains of RAN1, models of the ETR1-RAN1 interaction were created based on multimeric modeling of the proposed interactions of the individual MBDs with ETR1. Moreover, unambiguous restraints were employed to represent the loop between MBD1 and MBD2.

AlphaFold3 (Abramson *et al*., 2024) is supposed to have an improved prediction potential for protein-protein complexes. To verify whether an improvement in the complex prediction also applies to ETR1/RAN1, the ETR1/RAN1 complex was remodeled (Figure S2). Models 1 to 3 show interaction models that are somewhat structurally related (RMSD < 10 Å). All MBDs are located less than 5 Å away from ETR1, with MBD1 positioned on the opposite side of ETR1 relative to MBD2 and MBD3. This is in contrast with model 4, in which the MBDs are all positioned on the same side of ETR1. All MBDs remain in the proximity of ETR1 within distances less than 5 Å. In model 5, none of the MBDs of RAN1 are directed towards ETR1. Based on these observations and the pLDDT and PAE values (Figure S2A, B), model 1 was further investigated.

The membrane plane orientation of ETR1 and RAN1 was predicted based on the structures of RAN1 (Uniprot: Q9S7J8) and the full-length ETR1 dimers employing the OPM3.0 server (Lomize *et al*., 2022). The membrane plane orientations were obtained for both the HADDOCK and AlphaFold3 models to verify whether both transmembrane domains are located in the same membrane plane (Figure 5 and Figure S2).

### Prediction of structural models of chaperone-chaperone interactions

Heterodimers of ATX1, CCHΔ, MBD1, MBD2 and MBD3 were predicted with ColabFold 1.5.2 (Mirdita *et al*., 2021). The individual monomers show the typical ATX1-like fold, and for ATX1, CCH, and MBD1 and 2, the cysteines of the copper-binding motif “MxCxxC” are oriented in a face-to-face conformation, as anticipated from homologous structures in the Protein Data Bank (PDB) (Banci *et al*., 2006) (*Saccharomyces cerevisiae*, Atx1-Cu(I)-Ccc2a complex PDB ID: 2GGP) (Rodriguez-Granillo and Wittung-Stafshede, 2008) and required for direct copper transfer. In the case of MBD3, face-to-face but also face-to-back and back-to-back conformations were predicted. The matrix over the PAE (predicted alignment error) indicates the high accuracy of the monomer and dimer models (Figure 4). As the complexes generated by ColabFold lack the linker connecting MBD1 and MBD2, which is of approximately eight residues in length, MODELLER 10.5 (Eswar *et al*., 2006) was used to generate a complete model (Figure 4E).

### Multiple sequence alignment

The amino acid sequences of the copper chaperones ATX1 and CCH as well as the metal-binding domains of P-type ATPases HMA5 and HMA7 (RAN1) from *Arabidopsis thaliana* were obtained from the UniProt database (Consortium, 2021). Amino acid sequences were subjected to a multiple sequence alignment using the Clustal Omega (Madeira *et al*., 2022) webserver. Alignments were visualized using Jalview version 2.11.1.3 (Waterhouse *et al*., 2009) and colored according to the Clustal X color scheme. Identifiers for the proteins/domains of proteins used for the multiple sequence alignment are: ATX1 (Q94BT9), CCH (O82089), HMA5 (Q9SH30), and HMA7 (RAN1) (Q9S7J8). Amino acid sequences of CCH, HMA5, and HMA7 were trimmed to encompass only the sequences of their metal binding domains, as indicated in the alignment. Sequences of MBD1-3 of HMA7 represent the amino acid sequences of the proteins used for the *in vitro* experiments.

### Cloning

Plasmids for the heterologous expression of MBD1, MBD2, and MBD3 were constructed by inserting the corresponding coding DNA sequence into the pETEV16b plasmid backbone using the *Gibson*-assembly (Gibson *et al*., 2009) method. The plasmid pETEV16b is derived from the pET-16b (Novagen) expression vector and contains a tobacco etch virus (TEV) protease cleavage site instead of the Factor Xa cleavage site present in plasmid pET-16b. The DNA inserts were generated by polymerase chain reaction (PCR) using primers listed in Table S3 and the expression vector pETEV16b-RAN1, a vector present in our laboratory containing the full-length coding sequence of *A. thaliana* RAN1 *(Hoppen et al., 2019)*, as a template. Polymerase chain reaction (PCR) products were generated using Q5 high-fidelity DNA polymerase (NEB), subjected to treatment with DpnI (NEB) to remove the methylated DNA template and purified using the Illustra™ GFX™ PCR DNA and gel band purification kit (Cytiva). The purified PCR products were used at a 3:1 ratio (insert:vector backbone) in the *Gibson*-assembly reaction to generate the final expression vectors. For this, a master mix containing DNA polymerase, exonuclease, and DNA ligase was added followed by incubation at 50°C for 60 min. DNA sequences were confirmed by *Sanger*-sequencing (Microsynth Seqlab).

### Expression and purification of recombinant proteins

The heterologous expression of MBDs was achieved by transforming *E. coli* BL21 Gold (DE3) cells (Stratagene) with the corresponding expression plasmid and selecting them on 1.5 % (w/v) 2YT-agar plates (5 g/l NaCl, 10 g/l yeast extract, 15 g/l tryptone, pH 7.3) containing 100 µg/ml ampicillin. The expression of ETR1_GAF^1-316^ was achieved using *E. coli* C41(DE3)ΔompFΔacrAB (Kanonenberg *et al*., 2019) (referred to as C41ΔΔ in the henceforth) and the expression vector pETEV16b_ETR1_GAF^1-316^ were used. A 500 ml overnight culture was grown in 2YT medium supplemented with 100 µg/ml ampicillin in a 1 l unbaffled Erlenmeyer flask at 30°C while shaking at 180 rpm. The overnight cultures were then used to inoculate expression cultures to a starting OD_600_ of 0.15. Expression cultures consisting of 500 ml 2YT-medium supplemented with 100 µg/ml ampicillin and 2 % (v/v) ethanol in 1 l baffled Erlenmeyer flasks were grown at 30°C while shaking at 180 rpm. The cultures for the expression of ETR1_GAF^1-316^ in *E. coli* C41ΔΔ did not contain ethanol. The temperature was reduced to 16°C as soon as the expression cultures reached an OD_600_ of 0.4. Protein expression was induced by the addition of isopropyl β-D-1-thiogalactopyranoside (IPTG) to a final concentration of 0.5 mM when cultures reached an OD_600_ between 0.6 and 0.8. Cells were further grown over night (17-20 h) and harvested by centrifugation at 7,000xg for 15 min and 4°C. They were then flash-frozen in liquid nitrogen and stored at -20°C until further use. Typically, 6 l of culture medium was used for a single expression.

For purification, cells were resuspended in a lysis buffer (see below) containing DNaseI (PanReac AppliChem) and lysed by passing twice through a cell disruption system (Constant Systems Limited) at 2 kbar and 5°C for MBDs and 2.4 kbar for ETR1_GAF^1-^_316_. For MBDs, lysis buffer A (300 mM NaCl, 50 mM HEPES, 1 mM DTT, 10 % (w/v) glycerol, 0.002 % (w/v) PMSF, pH 7.2) was used whereas lysis buffer B (140 mM NaCl, 2.7 mM KCl, 10 mM Na_2_HPO_4_, 1.8 mM KH_2_PO_4_, 10 % (w/v) glycerol, 0.002 % PMSF, pH 8.0) was used for ETR1_GAF^1-316^. In order to purify the membrane protein ETR1_GAF^1-316^, cell membranes were isolated first and membrane proteins were solubilized using detergent. Therefore, the cell lysate was centrifuged at 14,000xg for 30 min and the supernatant was subjected to a second centrifugation at 40,000xg for 30 min. The supernatant was discarded and membrane pellets were washed by resuspending in lysis buffer B. Finally, membranes were isolated by centrifugation at 34,000xg for 30 min, shock frozen in liquid nitrogen, and stored at -80°C until further use (Schott-Verdugo *et al*., 2019). ETR1_GAF^1-316^ was solubilized by resuspending membrane pellets at a final concentration of 30 mg/ml in solubilization buffer (200 mM NaCl, 50 mM Tris/HCl, 1.5 % (w/v) Fos-Choline-14, 0.002 % PMSF, pH 8.0) and gentle stirring for one hour at 4°C. The cell lysates of soluble MBDs or the solubilized membranes were then cleared by ultracentrifugation at 100,000xg for 1 h (MBDs) or 230,000xg for 30 min (ETR1_GAF^1-316^). Proteins were purified by immobilized metal ion affinity chromatography (IMAC) using an ÄKTAPrime plus system (GE Healthcare, Illinois, USA). In order to purify the MBDs, supernatants of the cleared cell lysates were loaded on two connected 5 ml HisTrap FF Ni-NTA columns (GE Healthcare) equilibrated in lysis buffer A. The column was washed with at least 30 column volumes (CV) of lysis buffer A followed by a second washing step of 15 CV using lysis buffer A + 100 mM imidazole to remove unspecifically bound proteins. MBDs were then eluted using lysis buffer A + 500 mM imidazole. Elution fractions were pooled, concentrated to a sample volume of 2.5 ml using a 3 kDa molecular weight cut-off (MWCO) Amicon^TM^ Ultra-15 centrifugal filter (Millipore) and buffer exchanged into lysis buffer A on a PD-10 desalting column (Cytiva). Protein concentration was determined based on the absorption at 280 nm and the corresponding molar extinction coefficient, which was calculated from the amino acid sequence using the ProtParam tool *(Gasteiger et al., 2005)*. Glycerol from an 80 % (w/v) stock solution was added to a final concentration of 20 % (w/v) followed by flash-freezing in liquid nitrogen and storage at -80°C. Homogeneity of proteins was achieved by size exclusion chromatography (SEC). For this, proteins were thawed on ice and a 500 µL sample was loaded onto a Superdex Increase^TM^ 75 (Cytiva) column equilibrated in degassed SEC buffer (300 mM NaCl, 50 mM HEPES, pH 7.2). The flow rate used for SEC was 0.2 ml/min. Elution fractions were pooled, concentrated using 0.5 mL centrifugal filters with a 3 kDa MWCO (Millipore), and prepared for storage as described above. For storage, samples were divided into 30-50 µl aliquots in PCR reaction tubes. All steps were carried out at 4°C. However, samples of MBD3 were concentrated at 10°C and thawed at room temperature due to the poor solubility of MBD3 at low temperatures. ETR1_GAF^1-316^ was purified in a similar way to the full-length receptor as described in (Milic *et al*., 2018) with some minor changes. In short, the supernatant from the centrifuged, solubilized membranes was loaded onto a 5 ml HisTrap FF Ni-NTA (GE Healthcare, Illinois, USA) that had been equilibrated in buffer A (200 mM NaCl, 50 mM Tris/HCl, 0.015 % (w/v) Fos-Choline-14, 0.002 % PMSF, pH 8.0). The column was then washed with 50 CV buffer A, 20 CV ATP buffer (buffer A + 50 mM KCl, 20 mM MgCl_2_, and 10 mM adenosine triphosphate (ATP)), again with 10 CV buffer A and 4 CV buffer A + 50 mM imidazole in order to remove unspecifically bound proteins. ETR1_GAF^1-316^ was eluted from the resin by the addition of buffer A + 250 mM imidazole, concentrated to a final volume of 2.5 ml using an Amicon^TM^ Ultra-15 centrifugal filter with a 10 kDa MWCO (Millipore) and used directly for labeling with AlexaFluor^TM^ 488-NHS (ThermoFisher Scientific). Copper chaperones ATX1 and CCH were expressed and purified as previously described in reference *(Hoppen et al., 2019)*.

### Labeling of recombinant proteins for microscale thermophoresis

Freshly purified ETR1_GAF^1-316^ was buffer exchanged into labeling buffer (300 mM NaCl, 50 mM K_2_HPO_4_/KH_2_PO_4_, 0.015 % (w/v) Fos-Choline-14, pH 8.0) using a PD-10 desalting column (Cytiva, Marlborough, USA) and concentrated to a volume of 2.5 ml as described above. A 10 mg/ml stock solution of the amine-reactive dye AlexaFluor^TM^ 488-NHS (ThermoFisher Scientific) was prepared in anhydrous dimethyl sulfoxide (DMSO). From this stock, the dye was added to the protein in a 2.5-fold molar excess and the reaction mixture was incubated on a rotary shaker at room temperature for 1 h in the dark. The reaction was then quenched by buffer exchange into storage buffer (300 mM NaCl, 50 mM Tris/HCl, 0.015 % (w/v) Fos-Choline-14, pH 8.0) using a PD-10 desalting column (Cytiva) to efficiently remove unreacted dye and labeling buffer components. The labeled protein was then subjected to centrifugation at 230,000xg and 4°C for 30 min. The supernatant was transferred to a fresh reaction tube and the degree of labeling (DOL) as well as the protein concentration were determined as described by the manufacturer. The sample was stored with 20 % (w/v) glycerol added and split into 50 µl aliquots. These aliquots were then flash-frozen in liquid nitrogen and stored at -80°C. Copper chaperones ATX1 and CCH were labeled as described in *(Hoppen et al., 2019)*. All labeled proteins had a DOL of between 50% and 100%.

### Microscale thermophoresis

AlexaFluor488^TM^-labeled ETR1_GAF^1-316^ was diluted in MST-F buffer (300 mM NaCl, 50 mM HEPES, 0.03 % (w/v) Fos-Choline-14, pH 7.2) to a concentration of 100 nM. Aliquots of MBDs were first diluted to a glycerol concentration of 10 % (w/v) using SEC buffer and then further diluted to 200 µM using MST-G buffer (300 mM NaCl, 50 mM HEPES, 10 % (w/v) glycerol, pH 7.2). First, serial dilution series of MBDs ranging from 200 µM to 48.8 nM were prepared in MST-G buffer. Next, equal volumes of the dilution series and labeled ETR1_GAF^1-316^ were mixed resulting in final concentrations of MBDs ranging from 100 µM to 24.4 nM and 50 nM of labeled ETR1_GAF^1-316^. Samples were then incubated for 10 min in the dark prior to loading into premium glass capillaries. Measurements were performed on a Monolith NT.115 (both NanoTemper Technologies) with the excitation power set to 50 % and the MST power set to 40 %. The same settings were used for interactions of labeled ATX1 and MBDs. However, for interaction studies involving labeled CCH, the LED power had to be adjusted to 80 % while the MST power remained at 40 %. AlexaFluor488^TM^-labeled copper chaperones ATX1 and CCH were pre-diluted in MST-T buffer (300 mM NaCl, 50 mM HEPES, 0.1 % (w/v) Tween20, pH 7.2) to a concentration of 200 nM and added to the dilution series of the individual MBDs. For microscale thermophoresis experiments investigating the interaction of ATX1 or CCH with MBDs, 1:2 dilution series of MBDs were prepared in MST-G buffer. Final concentrations of labeled ATX1 or CCH were 100 nM, while concentrations of MBD1 ranged from 300 µM to 0.19 nM. Final concentrations of MBD2 and MBD3 ranged from 100 µM to 0.19 nM (MBD2) and 0.06 nM (MBD3). Interactions of individual MBDs with labeled ETR1_GAF^1-316^ were analyzed using thermophoresis with T-jump as the evaluation strategy. For experiments involving labeled copper chaperones ATX1 or CCH, initial fluorescence was chosen for data analysis due to strong, concentration-dependent changes in the initial fluorescence at increasing MBD concentrations. For these samples, SD-tests were performed according to the manufacturer to exclude material loss as a cause for changes in initial fluorescence. Data were fit to a one-site binding model using the MO.Affinity Analysis software v2.1.5 (NanoTemper Technologies). All measurements were performed in triplicate at room temperature.

### Copper Binding of MBDs

The appropriate volume of SEC-purified MBDs was added to a final volume of 150 µl, with a 20-fold molar excess of DTT using SEC buffer for dilution. The sample was then incubated for one hour either on ice (MBD1 and MBD2) or at room temperature (MBD3) to fully reduce the protein. The DTT was then removed by applying 75 µl of sample onto Zeba^TM^ spin desalting columns (ThermoFisher Scientific) with a 7 kDa MWCO, equilibrated in buffer SEC, according to the protocol provided by the manufacturer. Elution fractions were pooled and further diluted to a final volume of 350 µl. The protein concentration was determined spectrophotometrically based on the absorption at 280 nm, as described above. Subsequently, a 1:1 dilution series consisting of 12 serial dilution steps was prepared using buffer SEC. The dilution series started at the concentration of the sample used to determine the protein concentration, and an additional sample of SEC buffer only was prepared. From the dilution series, 50 µl samples were transferred to a transparent flat-bottom 96-well microtiter plate (Sarsted) and mixed with an equal volume assay buffer (300 mM NaCl, 50 mM HEPES, 2 mM sodium ascorbate, 100 µM BCA, 40 µM CuCl, pH 7.5). Assay buffer was added last using a multichannel pipette and absorption at 562 nm was recorded on a Tecan infinite M200 Pro plate reader (Tecan) to monitor the transfer of copper from the chromophoric Cu(I)-BCA_2_-complex to the protein. Control samples probing for residual DTT were treated the same as the MBD samples but did not contain protein. In a further control experiment, a protein (MSP1E3) containing a deca-histidine tag was used to study the effect of the deca-histidine tag in the competition assay.

Measurements were performed in triplicate and all steps were carried out at room temperature unless stated otherwise. Data were fitted to a four-parameter logistic function with a shared Hill coefficient. In order to better identify differences in MBDs copper binding abilities, the Hill coefficient was adjusted by the fitting algorithm. Data were plotted and analyzed using OriginPro (Version 2021b, OriginLab Corporation).

The reaction of the purified MBD with BCA_2_-Cu(I) is described by the equilibrium

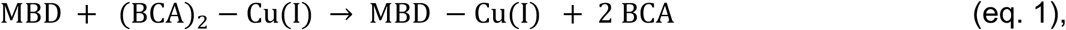

which consists of the two partial reactions

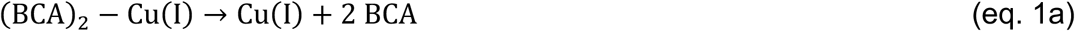

and

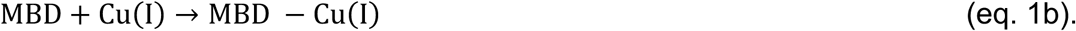

The individual equilibrium constants for eq. 1 and the total Cu(I) concentration are

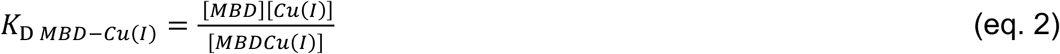

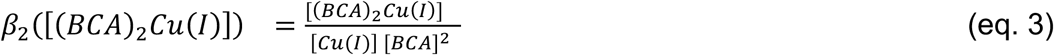

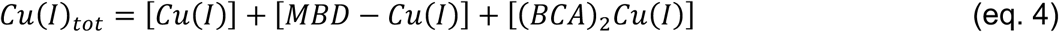

Using an approximation related to Cheng and Prusoff (Cheng and Prusoff, 1973), [MBDCu(I)] and [(BCA)_2_Cu(I)] from eq. 4 can be substituted with the corresponding equilibrium expressions (eqs. 2 and 3)

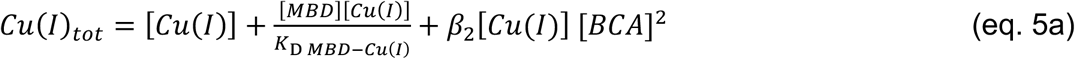

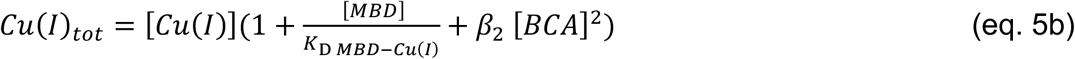

Replacing [Cu(I)] in eq. 3 with the expression from eq. 5b yields

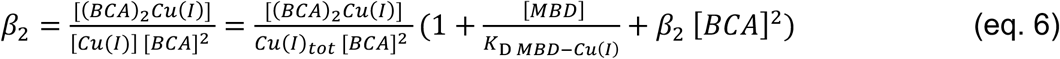

Rearranging to isolate [(BCA)_2_Cu(I)] yields

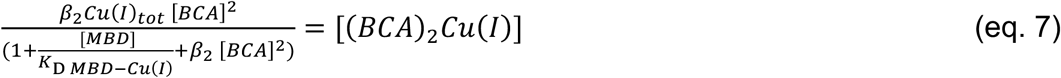

We now consider that the EC50 value is defined as the protein concentration at which the [BCA_2_Cu(I)] signal is half the signal when no protein is present. Note that due to the BCA excess in the experimental setup, the free BCA concentration is different when protein is present ([BCA_w/ protein_] or not [BCA_w/o protein_])

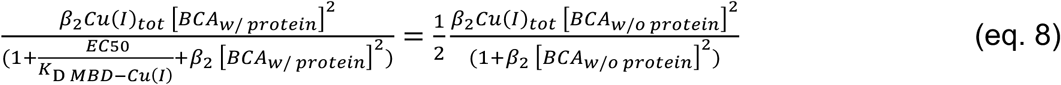

Thus, *K*_D MBD-Cu(I)_ can be calculated from the EC50 value, the respective BCA concentrations, and previous estimates of the formation constant of BCA_2_-Cu(I), β_2_ = 2 ⨉ 10^17^ M^-2^ (Xiao *et al*., 2008).

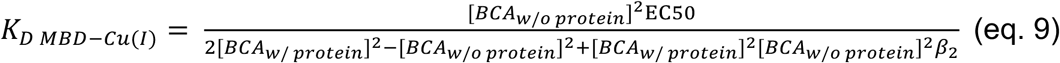

The EC50 values for the copper transfer from the chromophoric Cu(I)-BCA_2_ complex to MBDs 1 to 3 are given in Figure 2B.

[BCA_w/o protein_] was determined as the initial BCA concentration before protein addition, representing the BCA excess (10 ⨉ 10^-6^ M). [BCA_w/ protein_] was determined as the BCA concentration after protein addition at which the excess of BCA and 50% of the previously bound BCA are free, corresponding to a reduction of the BCA_2_Cu(I) signal by half as Cu binds to the protein (30 ⨉ 10^-6^ M).

This then leads to

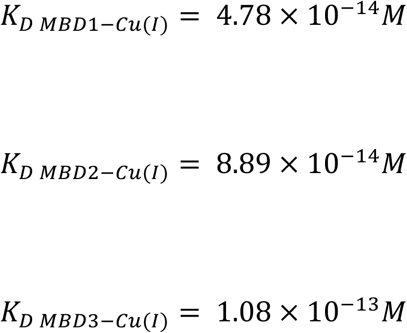

### Circular dichroism spectroscopy

Prior to the measurement, proteins were buffer exchanged into CD buffer (10 mM KCl, 10 mM K_2_HPO_4_/KH_2_PO_4_, pH 7.2) using Zeba^TM^ spin desalting columns (ThermoFisher Scientific) as described above and adjusted to a final concentration of 20 µM. Spectra were recorded from 190 nm to 320 nm on a Jasco J-815 spectropolarimeter (Jasco) at 20°C using a quartz glass cuvette with a path length of 1 mm. The scanning speed was 50 nm/min with a 1 nm bandwidth, the digital integration time (D.I.T) was 2 s and the step size was 0.1 or 1 nm. For each spectrum, eight measurements were accumulated. The BeStSel webserver (Micsonai *et al*., 2022; Micsonai *et al*., 2018; Micsonai *et al*., 2015) was used to estimate secondary structure contents from the obtained spectra, considering wavelengths in the range from 190 nm to 250 nm.

### Protein crystallization

For the crystallization of MBD3 of *A. thaliana* RAN1, aliquots of the protein purified by IMAC were thawed and further purified to homogeneity by SEC as described above. The purified protein was stored at 6-8°C overnight prior to crystallization. MBD3 was then crystallized using the sitting drop vapor diffusion method at 285.15 K (Hampel *et al*., 1968). Crystals were grown from a mixture containing 0.1 µL of protein solution (4.8 mg/mL protein, 50 mM HEPES pH 7.2, 300 mM NaCl) and 0.1 µL of precipitant (100 mM MES pH 6.5, 200 mM NaCl, 10% (w/v) PEG 4000). Rod-shaped crystals of approximately 300 ⨉ 20 µm formed within 75 days. Mineral oil (Sigma Aldrich, M3516) was applied to the crystallization drops before harvesting to prevent evaporation during handling and to serve as a cryo-protectant. The crystals were then harvested using lithographic loops (Dual Thickness MicroLoops LD, MiTeGen) and flash-cooled in liquid nitrogen. The X-ray diffraction data were collected at 100 K and a wavelength of 0.8856 Å at the European Synchrotron Radiation Facility (ESRF) beamline ID23-1 using an Eiger2 16M detector (Dectris). To minimize radiation damage, a helical data collection protocol was applied with an oscillation range of 0.1° per image, resulting in a dataset of 3600 images for a total oscillation width of 360°.

Data reduction, integration, and scaling were performed using the program DIALS (Winter *et al*., 2018). Based on data processing statistics and manual inspection of diffraction images, a subset of 1900 images (images 1250 to 3150) was used for final data processing. However, only 65% of all reflections were successfully indexed by DIALS when allowed to search for multiple lattices. This resulted in two distinct lattices, which were subsequently merged due to virtually identical unit cell parameters. The resolution cutoff was initially determined using a threshold for CC_½_ of 0.3 leading to a high-resolution cutoff of 1.78 Å. Due to low data completeness within the high-resolution shells caused by the chosen detector distance during data collection, this was eventually lowered to 1.98 Å.

Initial phases were obtained from molecular replacement (MR) using the AlphaFold2 (Jumper *et al*., 2021; Varadi *et al*., 2022) prediction of RAN1 as a template (AlphaFold DB: Q9S7J8). The model was cut to the boundaries of MBD3, and pLLDTs were converted to B-factors using tools provided by the PHENIX (Liebschner *et al*., 2019; Terwilliger *et al*., 2023; Terwilliger *et al*., 2022) software package before usage as a search model in PHASER (McCoy *et al*., 2007). The initial MR solution was used for automated model rebuilding using BUCCANEER (Cowtan, 2006) from the CCP4 software suite (Winn *et al*., 2011). The resulting model was then subjected to iterative manual model rebuilding using COOT (Emsley *et al*., 2010). Each iteration of manual rebuilding was followed by crystallographic refinement using PHENIX.refine (Afonine *et al*., 2012). Data collection and refinement statistics are summarized in Table S1. All figures in this study were prepared with PyMOL (The PyMOL Molecular Graphics System, Version 2.5 Schrödinger, LLC).

## Data availability

The atomic coordinates and structure factors were deposited in the Protein Data Bank (PDB, www.pdbe.org) under the accession code 8RNZ (DOI: 10.2210/pdb8rnz/pdb). Raw diffraction image data were deposited in the Zenodo repository (DOI: 10.5281/zenodo.10479001).

## Author contributions

G.G., D.D., H.G., and L.S.K conceived the project, D.D. and G.G designed and analyzed wet-lab experiments, L.S.K., S.S-V., and H.G. designed and analyzed computational experiments. D.D. performed wet-lab experiments, A.M. evaluated crystallographic data, L.S.K. and S.S-V. performed computational experiments. G.G. and H.G. secured funding and provided resources. All authors wrote and structured the manuscript.

## Acknowledgements

We are grateful for computational support and infrastructure provided by the “Zentrum für Informations-und Medientechnologie” (ZIM) at the Heinrich Heine University Düsseldorf and the computing time provided by the John von Neumann Institute for Computing (NIC) to HG on the supercomputer JUWELS at Jülich Supercomputing Centre (JSC) (user ID: VSK33, ETR1). We acknowledge the Center for Structural Studies (CSS) at the Heinrich Heine University Düsseldorf for assisting in screening protein crystallization conditions. The Center for Structural Studies is funded by the Deutsche Forschungsgemeinschaft (DFG Grant number 417919780). Further, we acknowledge the European Synchrotron Radiation Facility (ESRF) for provision of synchrotron radiation facilities under proposal number MX2485, and we would like to thank Alexander Popov for his assistance and support in using beamline ID23-1.

