## Supplementary Information for "Molecular Mechanism and Structural Models of Protein-Mediated Copper Transfer to the *Arabidopsis thaliana* Ethylene Receptor ETR1 at the ER Membrane"

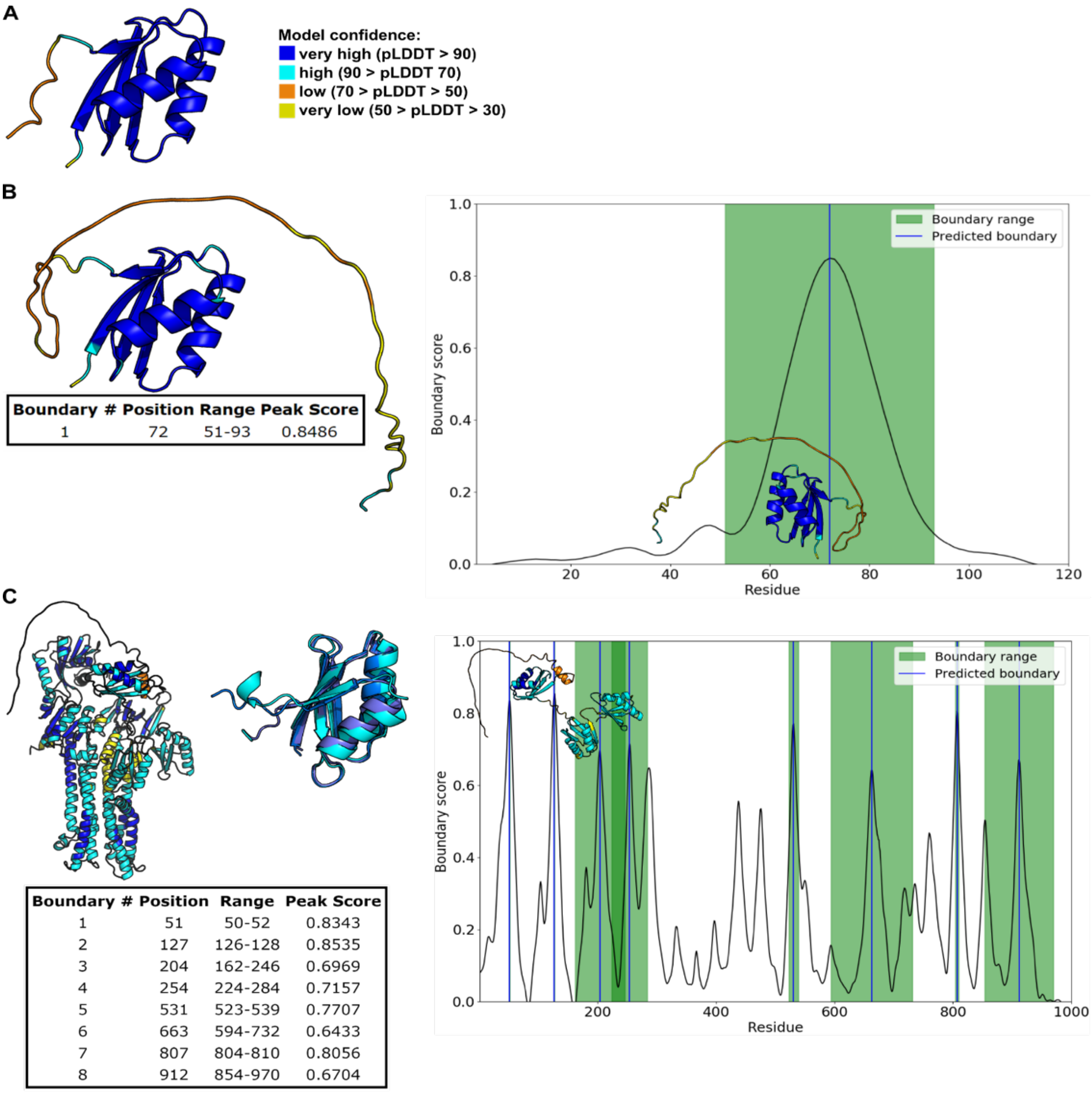


**Figure S1: Characterization of boundaries in CCH and RAN1 using AlphaFold2 and TopDomain. A:** Representation of the structural model of ATX1 generated with AlphaFold. Colors correspond to the pLDDT, with dark blue indicating pLDDT > 90. The model has the ATX1-like fold. Residues in secondary structure elements have high pLDDT values. **B:** Representation of the structural model of CCH generated with AlphaFold. Colors correspond to the pLDDT. The model has the ATX1-like fold and residues located in secondary structure elements have high pLDDT values. The pLDDT values begin to fall from residue 69. TopDomain predicts a boundary between residues 51-93, with the highest boundary score at residues 52-93. For CCH to be comparable to ATX1, the boundary to the unstructured domain was set at residue 76. **C:** Representation of the structural model of RAN1 generated by AlphaFold. Colors correspond to the pLDDT. The structure has ATX1-like folds at the N-terminus. Sequence parts with these folds have pLDDT > 70 and are linked by loops with lower pLDDT values. The transmembrane domains and topological domains have pLDDT values of 70-90. TopDomain predicts boundaries between residues 50 to 53, 126 to 128, 162 to 246, and 224 to 284. Since residue 54 still has a pLDDT < 70, the beginning of MBD1 was set to 55. The end of MBD1 was set to 130 to include the ATX1-like fold and also some of the residues with lower pLDDT as in the case of ATX1 and CCH. MBD2 starts with the first residue involved in the second ATX1-like fold according to the AlphaFold model and ends with residue 207, to include additionally some of the residues featuring a lower pLDDT as in the case of

MBD1. MBD3 starts with residue 205, as the linker between MBD2 and MBD3 is only four residues long, and ends with residue 283 to include additionally some of the following residues.

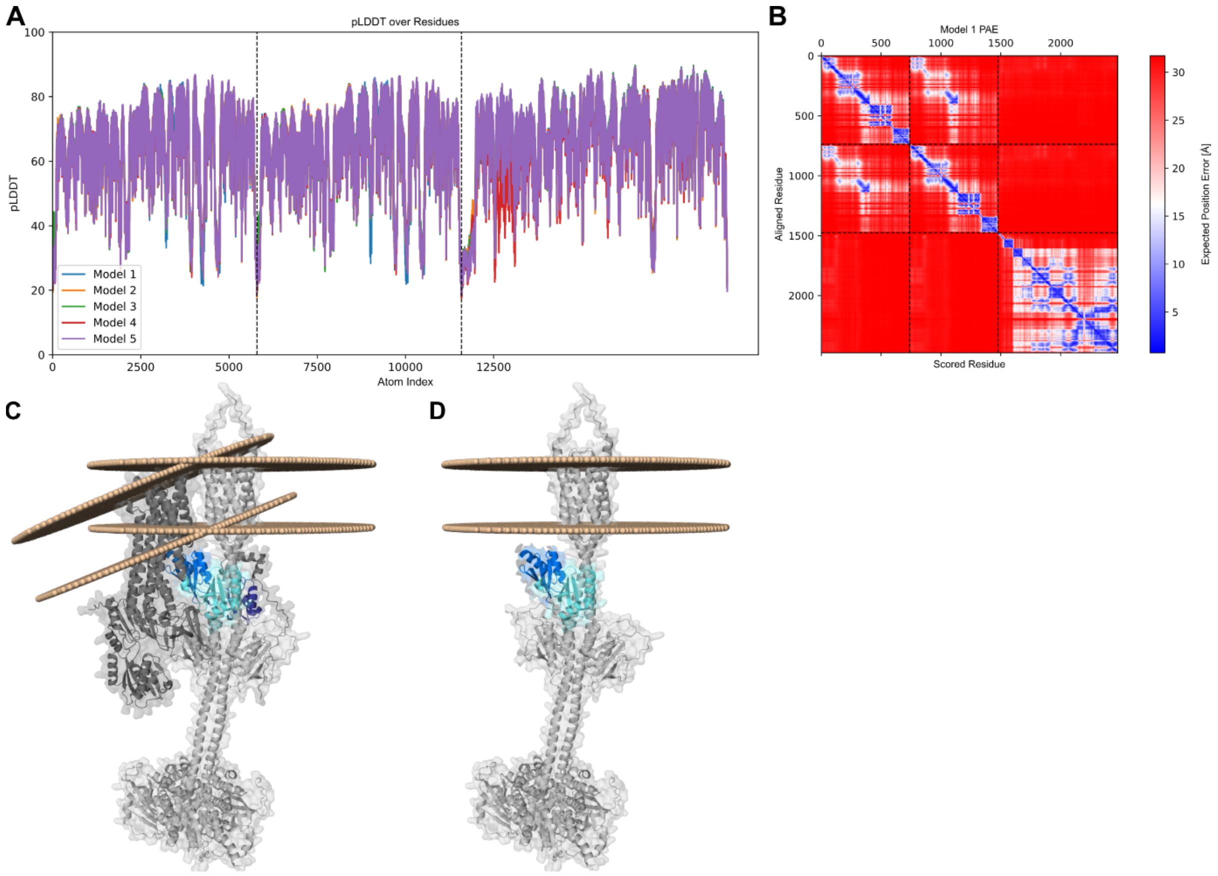


**Figure S2: RAN1/ETR1 complex predicted by AlphaFold3.** Visual inspection suggests that models 1 to 3 are somewhat structurally related (RMSD < 10 Å) and similar to the complexes created by protein-protein docking (Figure 5). In rank1-3, all MBDs feature distances of less than 5 Å to ETR1, where MBD1 is located on the opposite side of ETR1 as MBD2 and 3. **A:** pLDDT of the corresponding models. **B:** Predicted alignment error of model 1, also representative of models 2 and 3. **C:** Membrane orientation of RAN1 and ETR1. The TMDs of RAN1 and ETR1 would not be located in the same membrane. **D:** Predicted binding positions of MBD2 and MBD3 of RAN1. MBD2 and MBD3 bind to the same domains as in the docking approaches. The remaining domains of RAN1 are hidden for the sake of clarity.

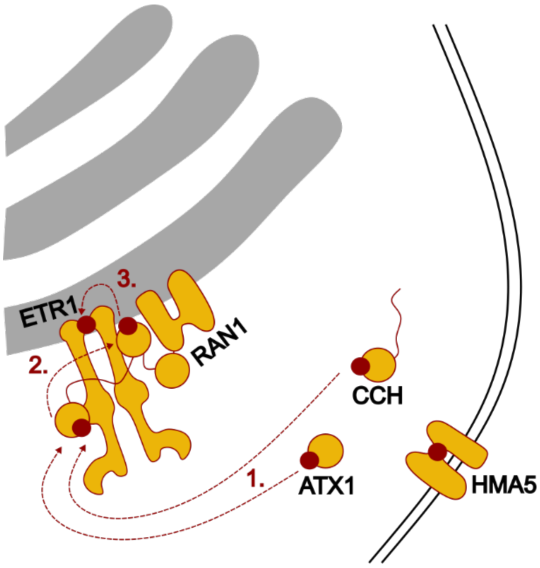


**Figure S3: Schematic representation of the copper relay hypothesis.** MBD1 accepts the copper from the soluble copper chaperones ATX1 and CCH and transfers the copper to MBD2. Due to MBD2's proximity to the ETR1 TMD, MBD2 finally transfers the copper to ETR1. MBD3 acts as an anchor stabilizing the ETR1/RAN1 complex.

**Table S1: Data collection and refinement statistics.** Statistics for the highest-resolution shell are shown in parentheses.

| | MBD3 (PDB-ID: 8RNZ) |
|---|---|
| **Data collection** | |
| Wavelength (Å) | 0.8856 |
| Resolution range (Å) | 51.48 – 1.98 (2.08 – 1.98) |
| Space group | $P2_12 2_1$ |
| Unit cell | |
| a, b, c (Å) | 50.755, 51.481, 62.722 |
| α, β, γ (°) | 90, 90, 90 |
| Total reflections | 80517 (11847) |
| Unique reflections | 11943 (1670) |
| Multiplicity | 6.7 (7.1) |
| Completeness (%) | 99.80 (99.24) |
| Mean $I/\sigma_I$ | 5.81 (0.74) |
| Wilson B-factor | 34.35 |
| $R_{merge}$ | 0.1204 (0.5585) |
| $R_{meas}$ | 0.1306 (0.6044) |
| $R_{pim}$ | 0.04984 (0.2279) |
| $CC_{½}$ | 0.996 (0.836) |
| CC* | 0.999 (0.954) |
| **Model and refinement** | |
| Reflections used in refinement | 11941 (1668) |
| Reflections used for $R_{free}$ | 568 (69) |
| $R_{work}$ | 0.1922 (0.3236) |
| $R_{free}$ | 0.2349 (0.3587) |
| Number of non-hydrogen atoms | 1309 |
| Macromolecules | 1229 |
| Ligands | 0 |
| Solvent | 80 |
| RMSD | |
| Bond lengths (Å) | 0.003 |
| Bond angles (°) | 0.540 |
| Ramachandran analysis | |
| Favored regions (%) | 100 |
| Allowed Regions (%) | 0 |
| Outliers (%) | 0 |
| Rotamer outliers (%) | 0 |
| Clashscore | 0.81 |
| Average B-Factor | 43.42 |
| Macromolecules | 43.34 |
| Solvent | 44.75 |

**Table S2: Secondary structure contents of MBDs 1 to 3 calculated from CD-spectra in the wavelength range from 190 nm to 250 nm.** Secondary structure contents were estimated by the BeStSel webserver, with the secondary structure elements $3_{10}$-helix, π-helix, bend, β-bridge, and loop/irregular regions being sorted into the category “others”.

| | **Secondary structure content [%]** | | | | **Spectral deviation** |
|---|---|---|---|---|---|
| | **α-helical** | **β-sheet** | **turn** | **others** | **NRMSD** |
| MBD1 | 12.8 | 29.5 | 12.6 | 45.1 | 0.009 |
| MBD2 | 8.8 | 32.4 | 13.6 | 45.2 | 0.010 |
| MBD3 | 20.4 | 24.4 | 10.4 | 44.8 | 0.008 |

**Table S3: Primers used for cloning of the metal binding domains (MBDs) 1 to 3 of RAN1 from *Arabidopsis thaliana*.**

| Primer | Sequence [5´-3´] |
|---|---|
| **ATXfld-1-F** | *ctgtattttcagggattaaggaagattcaggtcgga* |
| **ATXfld-1-R** | *cggatcctcgagttattgtgtctgttcttcagcc* |
| **ATXfld-2-F** | *ctgtattttcagggaactttggtgggtcaatttac* |
| **ATXfld-2-R** | *cggatcctcgagttacttatcctgctgattactctg* |
| **ATXfld-3-F** | *ctgtattttcagggacagcaggataagcttgttttaag* |
| **ATXfld-3-R** | *cggatcctcgagttaataagggctcataacacgc* |
| **pETEV16b-F** | *taactcgaggatccggctg* |
| **pTEV-R** | *tccctgaaaatacaggttttcatgg* |
